# Regenerative and non-regenerative transcriptional states of the human epicardium: from foetus to adult and back again

**DOI:** 10.1101/2021.10.11.462541

**Authors:** Vincent R. Knight-Schrijver, Hongorzul Davaapil, Alexander Ross, Xiaoling He, Ludovic Vallier, Laure Gambardella, Sanjay Sinha

## Abstract

Epicardial activation appears to be required for cardiac regeneration. Although reverting quiescent adult epicardium to an active neonatal or foetal state will likely represent a key therapeutic approach for human cardiac regeneration, the exact molecular differences between human adult and foetal epicardium are not understood. We used single-cell RNA sequencing to compare epicardial cells from both foetal and adult hearts. We found two foetal epicardial cell types, mesothelial and fibroblast-like, with only the mesothelial population present in adults. We also identified foetal-specific epicardial genes associated with regeneration and angiogenesis, and found that adult epicardium may be primed for immune and inflammatory responses. We predict that restoring the foetal epicardial state in human hearts would increase adult angiogenic potential. Finally, we demonstrated that human embryonic stem-cell derived epicardium is a valid model for the foetal epicardium and for investigating epicardial-mediated cardiac regeneration in humans. Our study defines regenerative programs in human foetal epicardium that are absent in the adult, brings human context to animal studies, and provides a roadmap for directing the epicardium in human heart regeneration.

## Introduction

A major challenge to human health is that the adult human heart does not regenerate; myocardial infarction (MI) causes a permanent non-contractile and non-conductive scar, which leads to chronic heart failure and arrhythmia. Much interest has followed the epicardium recently for its key role in heart development and potential to contribute to heart regeneration.

The epicardium emerges from the proepicardium during cardiogenesis as a mesothelial layer of cells surrounding the heart^1^. During development, epicardial cells can undergo epithelial to mesenchymal transition (EMT), resulting in the migration of epicardial-derived cells (EPDCs) into the myocardium^2^ where they may differentiate into smooth muscle cells, cardiac fibroblasts and a small proportion of endothelial cells^3–6^. Furthermore, developing epicardial cells and EPDCs secrete potent factors including WNT, FGFs and PDGFs which stimulate vasculogenesis and the proliferation and maturation of cells within the myocardial tissue^7^. These developmental abilities also translate into a regenerative role. Adult zebrafish hearts are capable of regeneration and developmental epicardial genes become highly expressed at the infarcted region, coinciding with the restoration of cardiac muscle^8,^. Likewise, when cardiac regeneration is seen in embryos and neonates of small and large mammals including humans^10–14^, the active epicardium responds with EMT and the secretion of angiogenic factors^15,16^. However, these studies also illustrate that any regenerative window in mammals soon disappears after birth.

In contrast, the adult mammalian epicardium is normally quiescent, with reduced secretory and migratory capacities and although it appears to reactivate after injury, the response may not be strong or rapid enough for sufficient regeneration^17^. However, there is evidence that a properly active epicardium can still regenerate adult mammalian hearts as studies have established the efficacy and essentiality of epicardial-directed repair mechanisms such as thymosin-*β* 4, FGFs, and even exosome-mediated signalling in successful cardiac regeneration^18–21^. Additionally, human embryonic stem cell (hESC)-derived epicardium (hESC-EPI) when administered alongside hESC-derived cardiomyocytes increases vascularisation, proliferation, and survival of myocardial tissue while reducing scar volume in infarcted rats^22^.

Altogether, the evidence suggests that the epicardium augments heart regeneration and that timely reactivation of epicardial programs offers a promising therapeutic strategy for treating MI in humans. However, without an in-depth understanding of the epicardium in humans, our ability to translate these models into a therapeutic context is limited. As it stands, it is still not fully known if the adult human epicardium retains gene expression from development or if its response to injury is similar to that seen in animals, or how it relates to hESC-EPI. Additionally, epicardial cells across many species are identified using WT1, TBX18 and TCF21^23,24^ but further divided into several subpopulations in zebrafish and in hESC-EPI^25,26^. However, this heterogeneity may not occur so clearly in mammals *in vivo*^27^ and it has not been fully explored in human tissues. We note though that adult human epicardium may be identified through its co-expression of BNC1 and MSLN^28^. Therefore, in light of this missing knowledge and we attempted to define the key factors of epicardial-derived regeneration that are lost in adults and aimed to capture the different transcriptional states of human epicardium, define age-associated changes in epicardial populations, and reveal distinct signalling pathways that are associated with foetal or adult epicardium.

We address our aims using single cell RNA sequencing (scRNAseq) to isolate epicardial cells *in silico* and mitigate biases from sorting and selection. Although scRNAseq data of both adult and foetal hearts have been generated and analysed independently, no attempt has been made to combine them^28,29^. For the first time, we integrated adult and foetal human hearts at a single cell resolution which allowed us to compare the epicardium in both stages. We approached our dataset from multiple angles and triangulated the epicardium in both adult and foetal cells using three levels of information. Firstly, we trained a classification model on a foetal sample to generate clustering-independent annotation of foetal and adult heart cells. We then expanded this individual cell resolution by clustering across all heart cells, grouping transcriptionally similar cell types together from both stages. Finally, we used annotations previously assigned to adult cells as a naive validation of adult and foetal cell alignment. This powerful approach converged on human epicardial cells spanning across human life and allowed us to i) identify foetal epicardial subtypes, ii) create a library of epicardial markers for translating animal studies, iii) reveal a regenerative epicardial signalling network not present in adult humans, and iv) validate hESC-EPIs as a model of the human foetal epicardium.

## Results

A total of 35,332 foetal cells were acquired from seven human foetal apices between the ages of 8 and 12 weeks post conception (Figure 1a). After quality control and preprocessing including the removal of erythrocytes, 14 054 foetal cells remained for further analysis (Supplementary Figure 1). In parallel, scRNAseq data covering 37,462 adult heart cells were obtained from the Heart Cell Atlas (HCA)^28^ selecting donors with epicardial cells (D3-D7) (Figure 1b). After applying identical quality control thresholds, we acquired a matrix of 33,102 adult heart cell transcriptomes (Supplementary Table 1).

**Figure 1.**
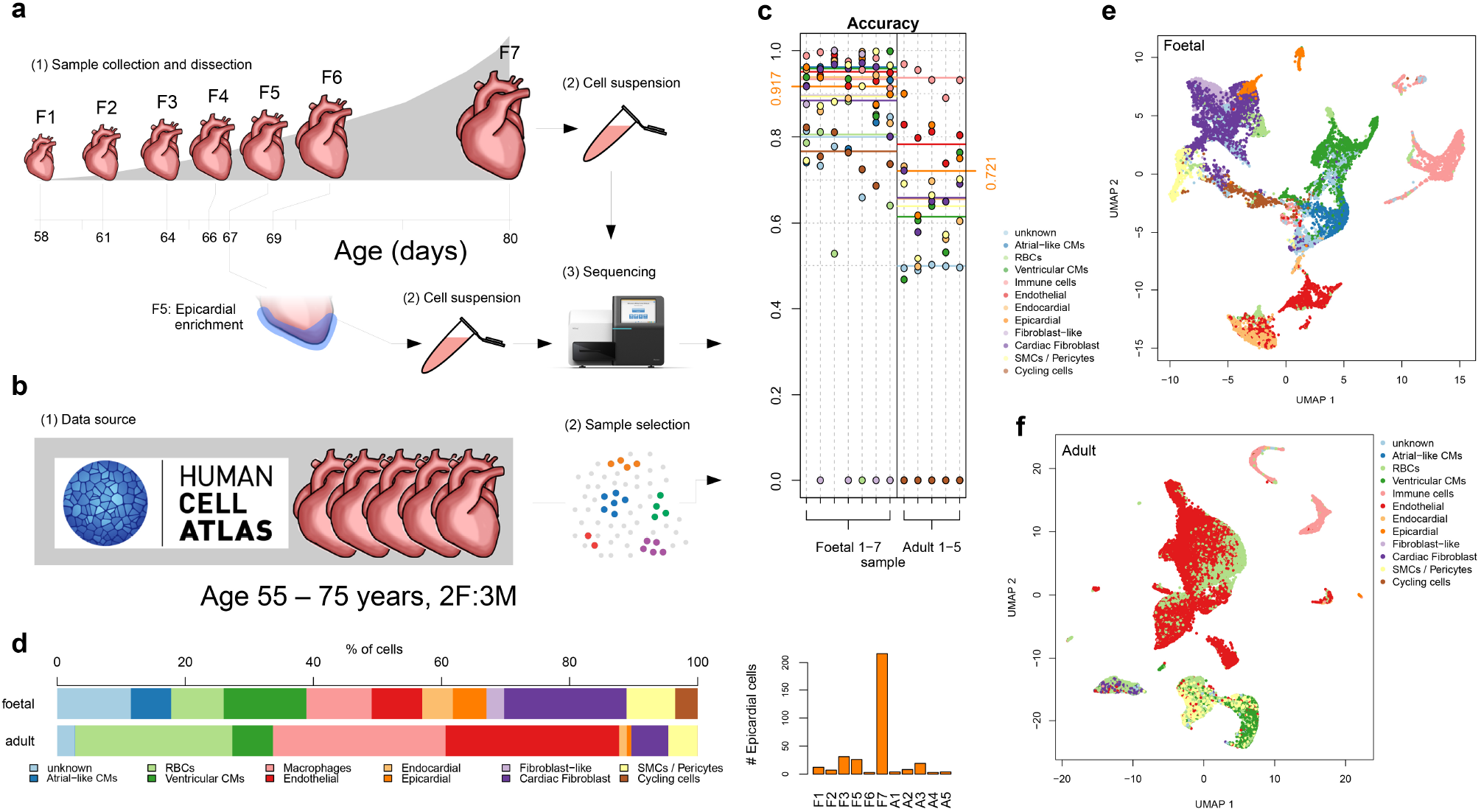
We predicted and compared the composition of adult and foetal hearts using a naïve Bayes classifier. **a**, Seven foetal heart samples between the ages of 8 and 12 weeks were dissociated into single cells and sequenced using the Illumina Chromium 10X platform. Epicardium was peeled in sampled F5; **b**, single cell RNA sequencing data of five adult hearts was acquired from the heart cell atlas; **c**, accuracy of the heart cell classifier in predicting cell types from all samples; **d**, predicted cell type composition of each stage and number of epicardial cells in each sample; **e**, UMAP projection of foetal data after 7 donors were integrated together; and **f**, UMAP projection of adult data after 5 donors were integrated together.

To extract epicardial cells and automate the annotation of cell types across all samples, we generated a heart cell classifier (HCC) on our first collected sample (sample F5). Briefly, we characterised 12 different cardiac cell types using density-based clustering (Supplementary Figure 2) and trained a naïve Bayes classifier (NBC) with an overall accuracy of over 95 % across all tissues and 99.9 % for epicardial cells (6-fold cross validation) (Supplementary Table 2). However, accuracy of the HCC was lower when applied independently to all foetal datasets (91.7 % compared with independent clustering) (Figure 1c, Supplementary Figure 1). In adult samples, the cluster containing *mesothelial* cells was classified as epicardium with a modest overall accuracy of 72.1 %, a low recall of 44 % but a high precision of 100 % (Supplementary Figure 2e). This precision suggest that while transcriptional states in the human epicardium clearly change with age, a specific epicardial signature remains conserved.

We attempted to enrich for epicardium in sample F5 by including an epicardial peeling with the dissociation. However, we found similar epicardial cell numbers in the three following apex-only samples (F1, F3 & F6) suggesting that this was not effective (Figure 1d). Therefore, we continued using only the apex of the last three foetal donors (F2, F4 & F7). Classification of cells suggested different compositions between adult and foetal hearts. The largest cluster of adult heart cells was composed of endothelial cells while most cells in foetal hearts appeared to be mixed fibroblasts and fibroblast-like cells (Figure 1e and f). We also captured more cardiomyocytes in foetal hearts, which may reflect a selection against larger mature adult cardiomyocytes by the sequencing pipeline. There was no significant difference between the number of epicardial cells in adult or foetal hearts (mean = 30, p > 0.05, student’s t-test). However, the oldest foetal sample F7 was over-enriched for epicardium (Figure 1d). Finally, we integrated the donors within their stages using Seurat’s integration workflow^30^, maintaining separate adult and foetal datasets, finding that HCC predictions localised well on two-dimensional uniform manifold approximation and projection (UMAP) embeddings (Figure 1e & f). Importantly, we noticed that epicardial-predicted cells in adult samples were isolated in one small cluster (Figure 1f) whereas epicardial cells in foetal hearts appeared to be divided into two clusters, one adjoining the mixture of fibroblasts (Figure 1e). This suggested that two epicardial cell types were found in foetal but not adult hearts.

### Unified adult and foetal hearts reveal stage-dependent cell types and a foetal-specific epicardial population

We combined both stages of heart cell data using a stratified sampling strategy aimed at minimising within-cluster variance caused by uneven donor sampling and between-cluster variance caused by uneven cell-type sampling (Supplementary Figure 3). We then integrated the sampled cells and reduced the dimensions using principal component analysis (PCA) and UMAP resulting in a unified picture of human developing and mature hearts at single cell resolution (Figure 2ai). Firstly, we found that both foetal and adult cells corresponded across the UMAP and that cell type predictions using the HCC were consistent, suggesting that most clusters were shared between stages (Figure 2aii). We then independently clustered the cells for downstream comparisons using Louvain clustering, revealing 23 clusters of adult and foetal cells (Figure 2aiii). Interestingly, we continued to find two epicardial clusters (15 and 17) identified by bonafide epicardial markers UPK3B and KRT19 (Figure 2b) as well as other epicardial markers such as ITLN1 and MSLN (Supplementary Tables 3 & 4). However, while cells in cluster 15 were sourced from both adult and foetal samples, cluster 17 was found to be 96 % foetal, suggesting a foetal epicardial cell type not present in the adult heart (Figure 3a). We noted that cluster 17 bridged the stage-shared epicardial cluster 15 and the aggregate of fibroblast clusters 1, 3, and 4 on the first two dimensions of the UMAP (Figure 2aii) and clearly expressed both epicardial and fibroblast genes (Figure 2b). This suggested that cluster 17 was a fibroblast-like epicardium possibly human mesenchymal EPDCs, while cluster 15 was mesothelial epicardium. Clusters 2, 4, and 13, were also over 90 % foetal, labelled as foetal cardiomyocytes, fibroblast-like cells, and cycling cells respectively. Interestingly, only cluster 20 was 100 % adult and contained stromal pericytes (Figure 2b, Supplementary Figure 3f).

**Figure 2.**
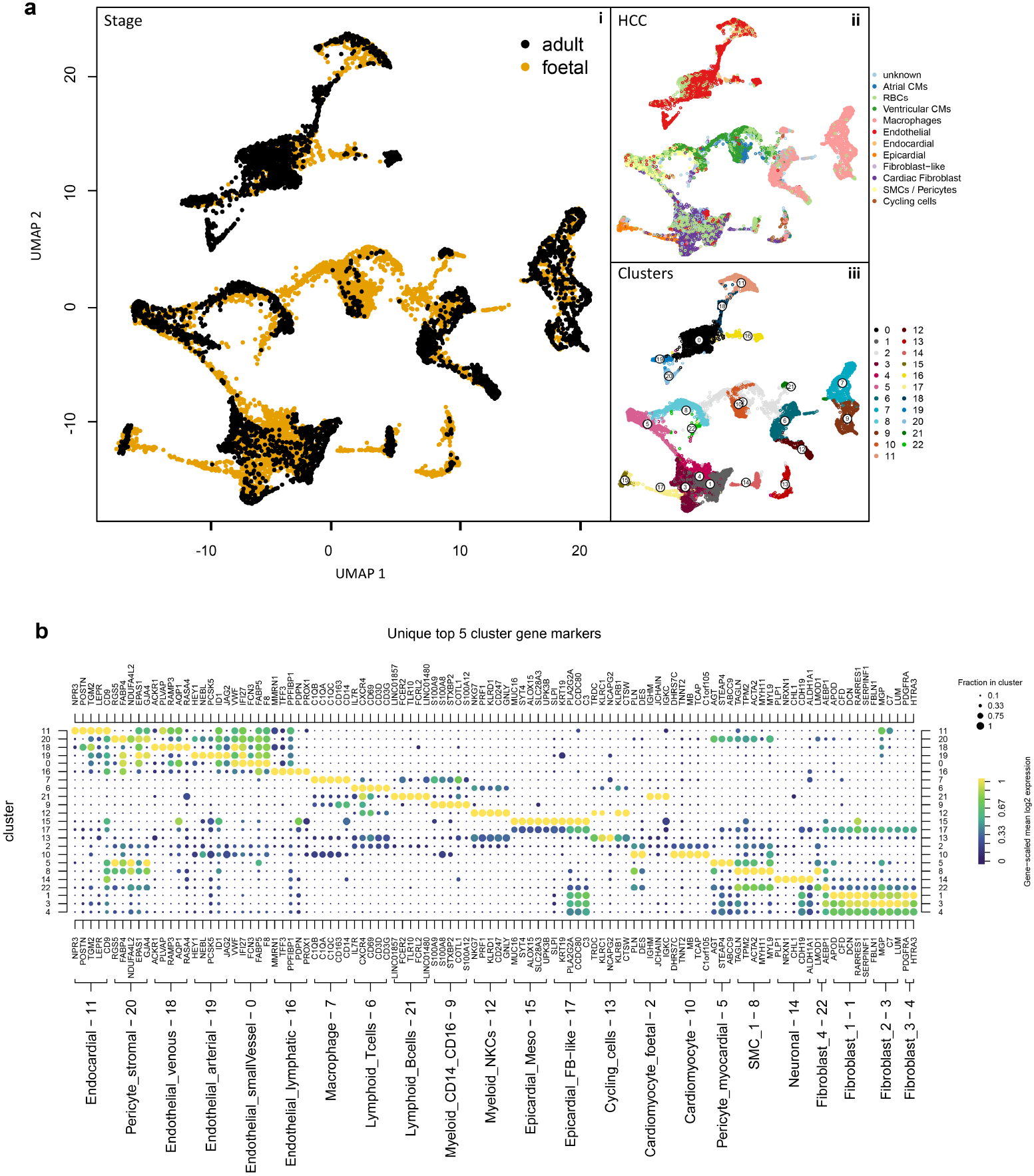
Adult and foetal samples were integrated using stratified sampling resulting in mostly corresponding cell type clusters but different epicardial populations. **a**, two-dimensional UMAP shows both foetal and adult heart cells integrated after a target sampling strategy and integration across 12 donors showing **i**, overlapping clusters, **ii**, heart cell classifier predictions for all cells, and **iii**, 23 stage-combined clusters identified using Louvain clustering. **b**, dot-plot of log2 fold-change and adjusted p values from Wilcoxon ranked sum tests illustrates top genes expressed in 23 stage-combined clusters with cluster 17 sharing epicardial and fibroblast markers.

**Figure 3.**
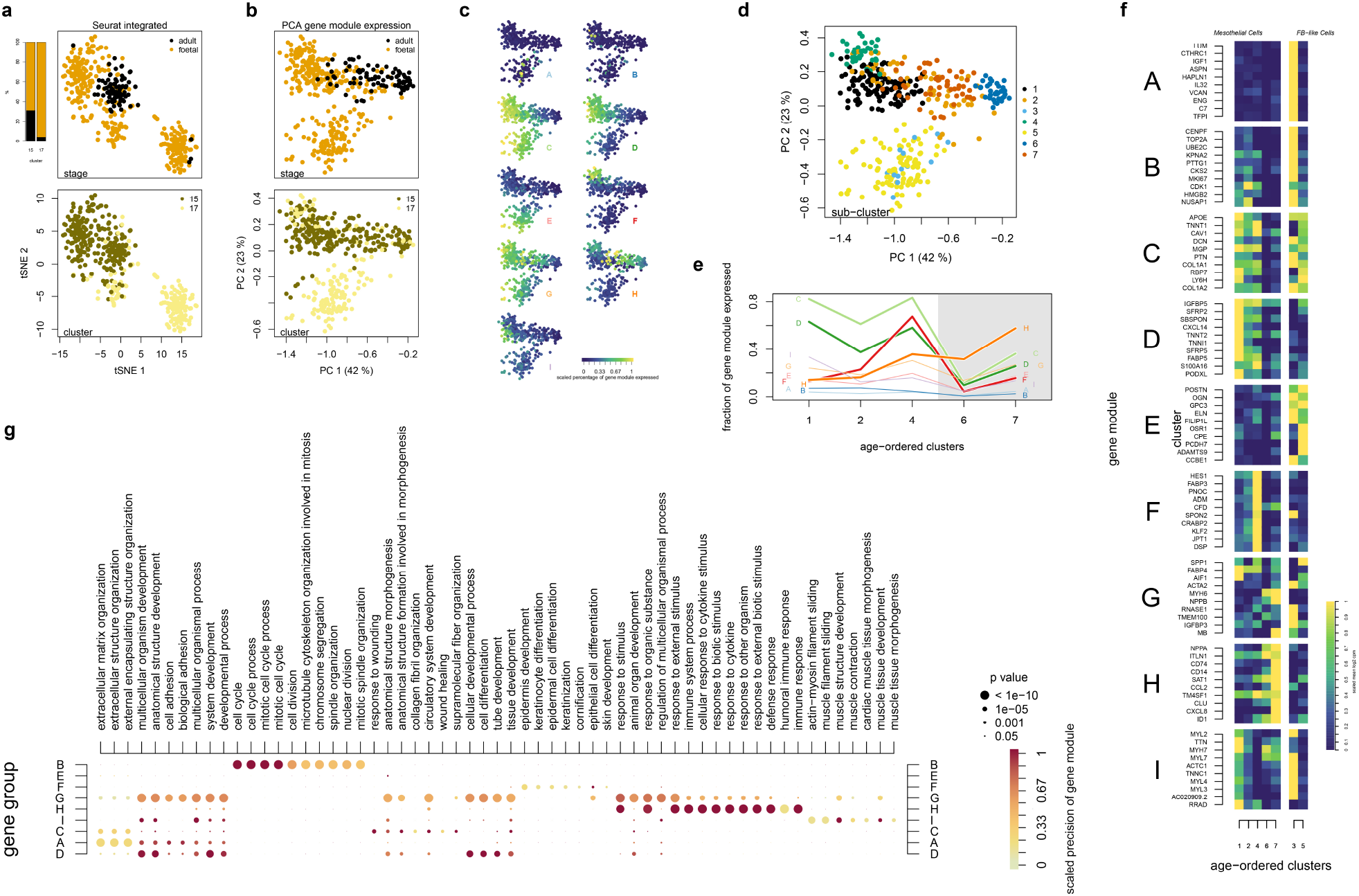
Foetal hearts contain a unique epicardial cell type as well as unique gene programs and functions, all reduced in the adult epicardium. **a**, a two-dimensional t-distributed stochastic neighbour-embedding plot of epicardial cells isolated from stage-integrated data and **b**, two dimensions of a principal component analysis of cells using their commitment to gene modules shows the separation of foetal-specific cell cluster 17 and the stage-shared cluster 15. **c** Epicardial cell commitment towards 9 gene modules across both stages projected onto principal components 1 and 2; **d**, 7 epicardial transcriptional states across both stages; **e** mean gene module commitment of epicardial cells ordered by the mean age of subclusters, the grey box shows adult epicardium; **f**, mean commitment of top 10 expression genes from each module; and **g**, a dot-plot summarising an over-representation analysis of modules displaying the top 10 gene ontology processes enriched in each gene module.

### EPDC gene programs are absent in adult human epicardium

We went on to further characterise the epicardial cell states by isolating clusters 15 and 17. During this process, outlying cells which were sparsely distributed across the entire foetal dataset were removed after finding a low number of UMIs, a high number of cells predicted as noise or an unknown cell type, and a low general epicardial marker expression (Supplementary Figure 4). Re-integration and clustering of the remaining epicardial cells more clearly separated mesothelial and fibroblast-like epicardial clusters 15 and 17 (Figure 3a). To assess the transcriptional programs present across these epicardial cells we focused on highly variable and epicardial-specific genes and sought to group co-expressed genes into distinct modules. This was achieved by clustering co-occurring genes together by their dropout patterns across our epicardial dataset using a binary representation of UMI counts. Then, significantly co-occurring genes were found using chi-square tests and the resulting statistics were clustered using the unsupervised Louvain method of community detection. This produced 9 unique gene expression modules characterising transcriptional states of the human epicardium (Figure 3c) (Supplementary Table 5). We used these modules as new epicardial variables by reducing the counts matrix to a gene module commitment matrix followed by a PCA (Figure 3b). We found that the first two principal components appeared to separate sample ages and mesothelial and fibroblast-like epicardium respectively (Figure 3b). Lastly, we clustered these cells according to module commitment and found 7 new epicardial transcriptional state clusters (Figure 3d). Notably, states 3 and 5 were part of the fibroblast-like epicardium outlined by their high commitment to gene module E (Figure 3c & f). This module includes the genes GPC3 (glypican 3) and POSTN (periostin) as further evidence of a human EPDC signature^24,31^ (Supplementary Figure 6). We also found high expression of COL3A1 in clusters 3 and 5 as seen in animal EPDCs (Supplementary Figure 6). Finally, gene module E was not found in adult cells suggesting that adult epicardial cells are not committed to this EPDC gene program and that these cells are not overly present in adult hearts.

### Human foetal epicardium is poised for regeneration but ages towards immune response

We assessed how epicardial cells change their commitment to these gene modules between stages by ranking the mesothelial clusters (1,2,4,6, & 7) by their mean age (Figure 3e). Firstly, we noticed that two gene modules (C and D) were markedly reduced in adult epicardial cells. On average, developing epicardial cells expressed 80 % of the genes clustered in pan-epicardial module C which decreased to between 10 % and 35 % in adult epicardium. We carried out an over-representation analysis of gene ontology biological processes using GprofileR. Among expected developmental terms, module C was significantly associated with regeneration including *response to wounding*, *wound healing*, and *circulatory system development* (Figure 3g) as well as *blood vessel development* and *canonical wnt signalling pathway* (Supplementary Figure 4d). A similar pattern was seen in module D, however unlike module C, D was only expressed by putative mesothelial cells (Figure 3f). Module D genes were more associated with *cell adhesion*, *cell differentiation* (Figure 3g) and *cell motility* (Supplementary Figure 4d). This suggests that movement and migration is highly regulated in foetal mesothelial states.

Secondly we found that the epicardium may be transitioning into a more immune-responsive state with age with increasing commitment to gene module H (Figure 3e & f). This adult mesothelial module was rich in associations with immune-related GO processes including *response to external stimulus*, *immune system process*, *cellular response to cytokine stimulus* (Figure 3g), and other broad terms such as *immune response* and *response to stress* (Supplementary Figure 4d). These terms were also partially shared by the pan-epicardial module G (Figure 3g). This unexpected result indicates an unexplored characteristic of epicardial ageing, which may be important for cardiovascular regeneration. A related observation was seen in a pseudo-bulk sampling correlation analysis showing that immune cells from both stages consistently clustered together while other heart cells clustered primarily within stages (Supplementary Figure 5). These findings suggested an altered epicardial immune-responsiveness in the absence of a substantially altered immune cell profile.

Other notable time-correlated gene modules were F and I, both seemingly associated with mesothelial development. On the one hand, module F appeared to increase with developmental time but failed to persist into adulthood, peaking at cluster 4 in cells belonging to the oldest foetal sample F7 (Figure 3e & f). This module was enriched for processes of *skin development*, *keratinization*, and *epidermis development*. On the other hand, gene module I decreased with development from its highest expression in the youngest foetal cells and was associated with *muscle development* and *muscle contraction*. Lastly, we ordered the epicardial cells by the first principal component to observe the transient expression of these age-associated genes (Supplementary Figure 6). Altogether, these results show an age-associated switch in epicardial commitment to gene programs moving away from important aspects of epicardial-regeneration such as paracrine signalling and migration.

### A library of precise epicardial markers differentiates between adult, foetal and fibroblast-like epicardium and reveals a WNT signature absent in adult epicardium

We then evaluated epicardial specific genes as markers of adult or foetal epicardium. This was achieved by carrying out differential expression analyses between the 23 heart cell-type clusters within each stage separately. In focusing on the mesothelial epicardium in cluster 15 we found 142 and 97 differentially upregulated genes in hearts from foetal or adult stages respectively and 57 upregulated genes that were shared (Wilcoxon rank-sum; adjusted p value < 1 × 10^−10^, log 2 fold-change > 0.5) (Figure 4a, Supplementary Table 6). We noted contrasting fold-change and expressions of top markers between adult and foetal epicardium which confirmed the stage-selectivity of these genes (Figure 4a & b). We then ranked epicardial markers from foetal, adult and shared gene sets by their ability to detect epicardial cells using precision, recall and F score.

**Figure 4.**
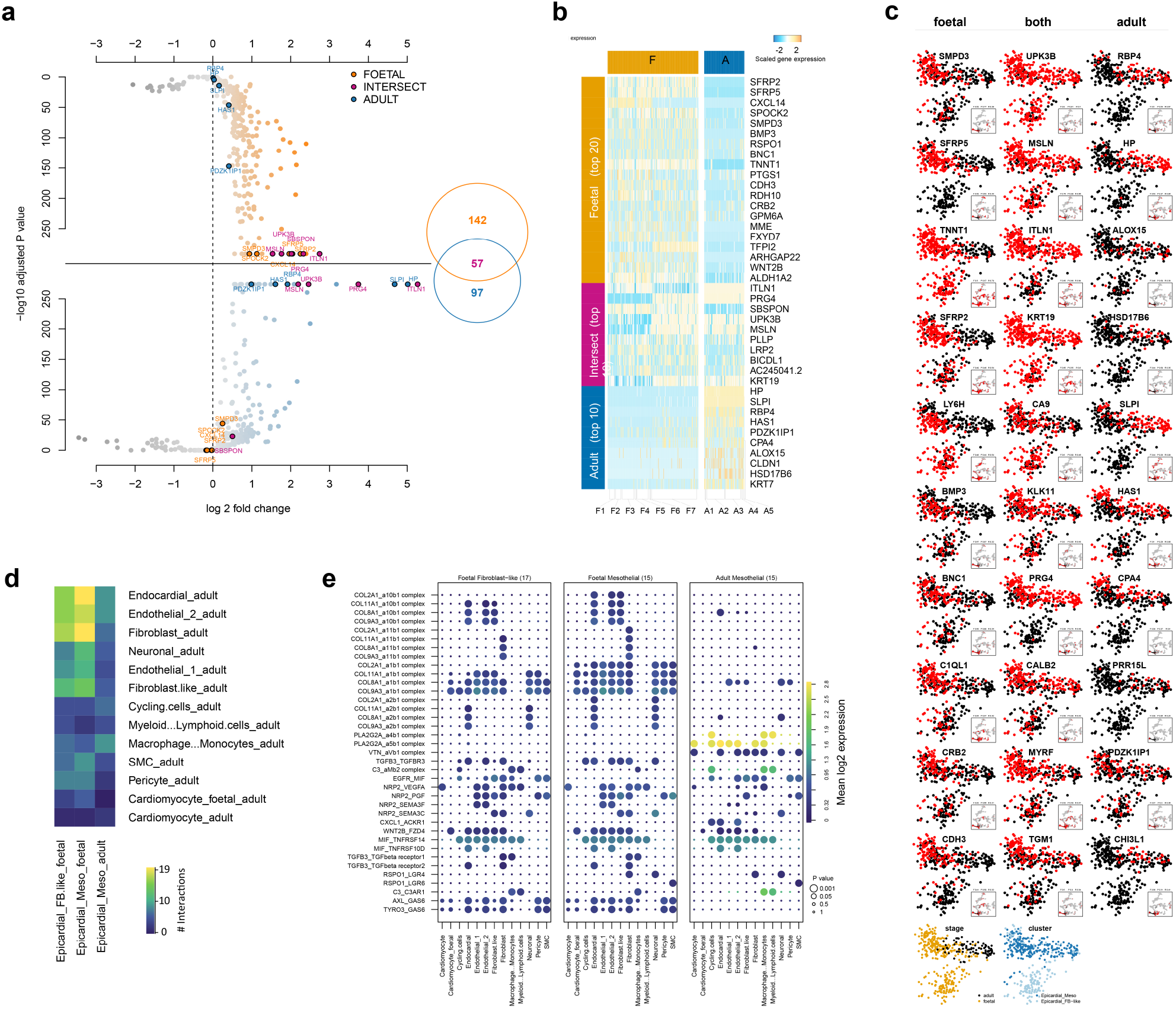
Foetal epicardial WNT genes, markers, and volume of communication are reduced in adult epicardium. Parallel Wilcoxon ranked sum tests carried out on cells within each stage yielded foetal and adult markers (p > 1e-10, log2 fold change > 0.5) shown in **a**, mirrored volcano plots labelled with the top 5 upregulated genes shared between and specific for each stage, **b**, a heatmap of top 20, 10, and 10 markers ordered by p value specific for foetal, both stages, and adult epicardium. Putative performance of genes as predictive markers is shown in **c**, a principal component analysis from gene module analysis binary data coloured as either expressed (red) or not expressed (black). The predictive performance is indicated in the subplot of each figure showing F score (F), precision (P), and recall (R) of identified markers with respect to other heart cells depicted in the sub plot. CellPhoneDB analysis plotted as **d**, heatmap of the number of interactions from epicardial clusters towards adult heart cell types; and **e**, dot-plot depicting the mean expression of both ligand and receptor, and significance of interaction.

Irrespective of age, UPK3B was the strongest predictive marker (Figure 4c). We also found the genes (MSLN, ITLN1, KRT19 and CA9), which confirmed their status as established epicardial markers and validated our library. However, we revealed novel markers such as TGM1 (transglutaminase) and CALB2 (calbindin 2) both highly specific for all epicardial cells with precisions of 0.88 and 0.9 respectively (Figure 4c). We noted that ITLN1 and MSLN were associated with the adult gene module H, suggesting that both ITLN1 and MSLN increase with epicardium maturation. ITLN1 was also expressed more frequently in mesothelial cells compared with fibroblast-like epicardial cells (Figure 4c). Interestingly, this property was shared by another predictive gene PRG4 (proteoglycan 4), which encodes for the protein *lubricin* found in pericardial fluid^32^. This selectivity further implied that cluster 17 cells were not on the surface of the heart. In adult hearts, we found that RBP4 (retinol-binding protein 4) was the best predictive marker of adult epicardium with a precision, recall and F score of 0.54, 0.67 and 0.6 respectively (Figure 4c). This was followed by HP (Haptoglobin), ALOX15 (arachidonate 15-lipoxygenase), HSD17B6 (hydroxysteroid 17-beta dehydrogenase 6), and SLPI (secretory leukocyte peptidase inhibitor) (Figure 4c). HP has previously been found in older human foetal epicardium^33^ and may be a marker of maturing mesothelial cells that persists into adulthood (Figure 4c). We found PRR15L (Proline Rich 15 Like) to be a highly precise marker of adult epicardium but with reduced sensitivity (precision = 0.95, recall = 0.24). PRR15L has been found in animal epicardium and may be useful as a translational marker alongside C3 (complement 3), DMKN (Dermokine) and GPM6A (Glycoprotein M6A) seen here and also previously found in mouse epicardium^34,35^.

In foetal epicardium, the most reliable marker was SMPD3 (sphingomyelin phosphodiesterase 3) with a high precision, recall and F-score (0.77, 0.39 & 0.52) followed by SFRP5 (secreted frizzled-related protein 2), TNNT1 (troponin t1, slow skeletal type), SFRP2, and LY6H (lymphocyte antigen 6 family member H) (Figure 4c). SPMD3 may be involved in packaging regenerative microRNAs for release into infarcted tissues^36^. TNNT1 had only recently been associated with epicardium^37^ and here it was not present in adults, while LY6H previously seen only in the brain and various tumours, is largely uncharacterised^38^. Additional genes of interest with high precision for foetal epicardium were BMP3 (bone morphogenetic protein 3) and RSPO1 (respondin-1) with precisions of 0.87 and 0.73 respectively (Figure 4c). SFRP5 also offered substantial predictive power for foetal epicardium while being confined to only mesothelial cells and could be used to distinguish EPDCs. Interestingly, many of these selectively foetal genes were WNT-related including SFRP2, SFRP5, RSPO1 and WNT2B. Finally, while we evaluated epicardial genes here for a predictive library of markers we also found typical epicardial genes BNC1 (basonuclin 1), ALDH1A2 (aldehyde dehydrogenase 1 family member A2) and PODXL (podocalyxin like) among foetal epicardial markers, suggesting a loss of well-established as well as novel regenerative epicardial functions with age.

### Restoring foetal epicardial states may increase communication, promoting regeneration and angiogenesis

Reduced cardiac regeneration in adults may be caused partly by a reduction in epicardial communication. Therefore, we sought to predict whether restoring foetal epicardial activity would drive regenerative communication in adult hearts. We selected secreted gene products from the stage-specific markers (Figure 4a & b) and predicted cell type communication coming from epicardium using CellPhoneDB. Firstly, we predicted that when compared with adult epicardium, the foetal epicardium interact more with the adult heart. Most communication was with endocardial, fibroblast, and endothelial populations with 19, 19, and 17 interactions respectively from foetal epicardium compared with 7, 5, and 7 interactions from adult epicardium. Agreeing with regenerative terms enriched in gene module C, we predicted that this volume of epicardial communication directed towards adult endocardial and endothelial cells could be pro-angiogenic. Signalling from foetal epicardium consisted of NRP2-mediated signalling via endothelial or immune cell derived VEGFA, and WNT2B communication between the foetal epicardium and FZD4 found on endothelial, endocardial, fibroblast and foetal / immature cardiomyocyte clusters. Reduced FZD4 activity has been seen to markedly decrease vascular density in kidneys^39^ while NRP2 and VEGFA is a well established route of angiogenesis. However, it was unknown whether this NRP2 was in soluble or membrane-anchored isoforms^40,41^. We also found epicardial RSPO1 in an interaction with LRG6 in smooth muscle cells, and with LRG4 in fibroblasts and neuronal cells but no expression of its usual receptor kremen 1. We also found TGFB3 signalling uniquely from foetal epicardium but not adult epicardium, agreeing with previous studies on its expression^42^. While TGFB proteins have a documented role in wound healing and fibrosis, TGFB3, normally low in adults^43^, has been seen at high levels after MI and might reduce scarring post-injury^44,45^. Additionally, TGFB3 was seen to decrease in a model of reduced EMT in mouse epicardium, coinciding with a reduction in vascular maturity of the mouse myocardium^31^.

Interestingly, we found that most foetal epicardial interactions were by collagens (Figure 4e). In particular, we noticed that collagens COL2A1 and COL11A1 were not expressed in other foetal tissues (Wilcoxon ranked sum test; adjusted P value < 1 × 10^−10^, log 2 fold-change > 0.5) and may be an epicardial component of matrix organisation. Unexpectedly, foetal communication with cardiomyocytes was low from both adult and foetal epicardium with only epicardial-sourced NRP2 interacting with cardiomyocyte VEGFA. From adult epicardium, a significant interaction between PLA2G2A and VTN was seen with cardiomyocyte integrins a5b1 / avb1 (Figure 4e).Interestingly, PLA2G2A has previously been associated with coronary heart disease and infarction^46^. These results provide evidence that *in situ* reactivation of foetal epicardial programs might increase regenerative communication with endothelial cells.

### hESC-derived epicardium models in vivo epicardium and expresses regenerative foetal genes

We previously harnessed active epicardial signalling to augment heart regeneration using hESC-EPI cells *in situ*^22^. However, the mechanism governing this success was unknown. To address this and identify commonalities between *in vitro* and *in vivo* epicardium, we harvested hESC-EPI in a scRNAseq timecourse during the last 9 days of their differentiation^47^ (Figure 5a). This protocol yields a heterogeneous two-population epicardium^26^ confirmed in our results as a divergent differentiation into two branches expressing either PODXL and BNC1 (lineage A), or TCF21 and THY1 (lineage B) (Figure 5b). We classified these cells using the *in vivo* HCC and found that epicardial predictions increased over differentiation, solely occurring in the PODXL+ branch (Figure 5e). In contrast, the TCF21+ branch was classified as fibroblasts (Figure 5e). This classification reflects the two epicardial populations 15 and 17 found*in vivo*.

**Figure 5.**
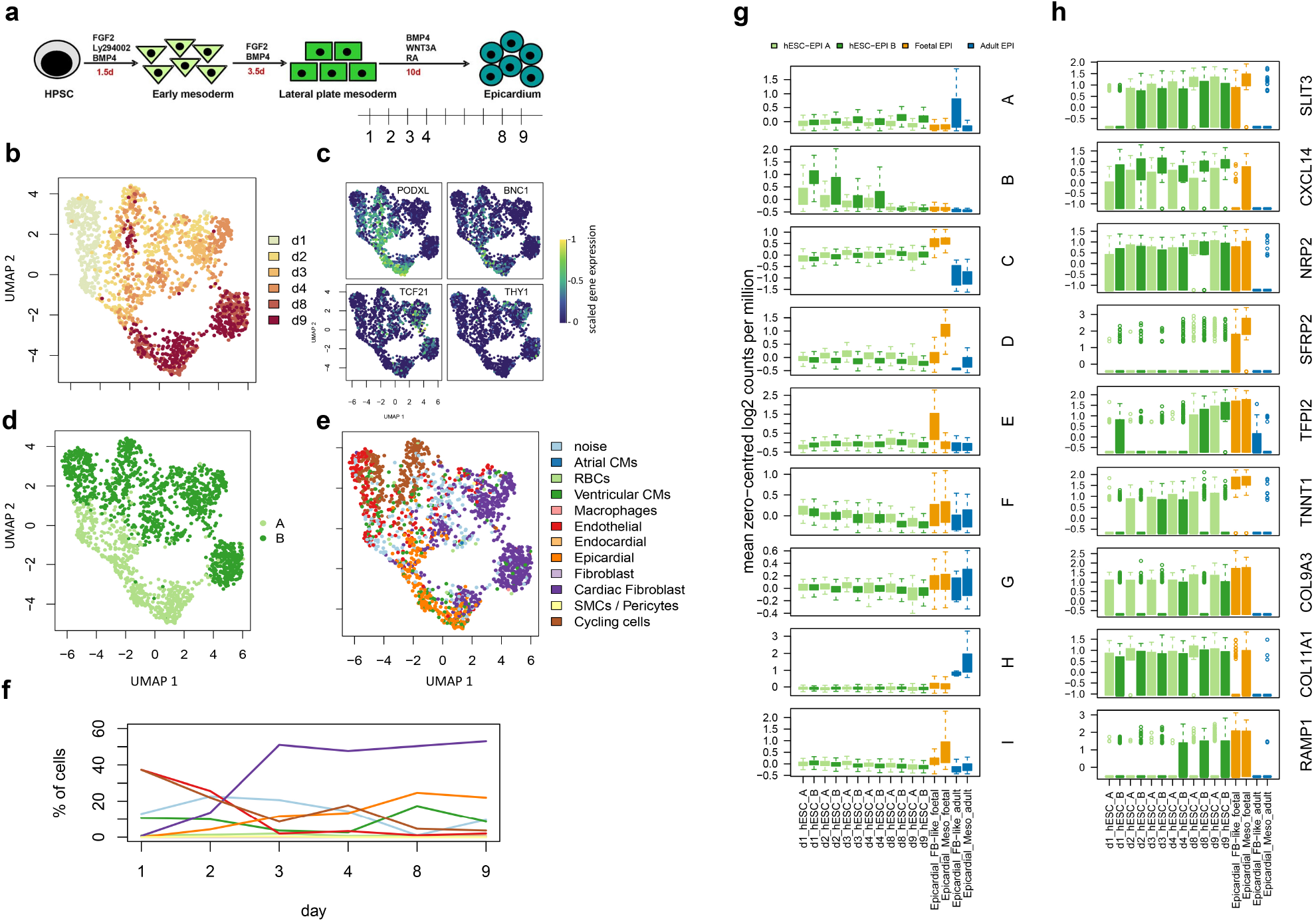
Transcriptional states of human embryonic-stem cell derived epicardium were compared with with foetal and adult epicardium. **a** 6 samples were taken between day 1 and day 9 after induction of epicardium from mesoderm stage using the differentiation protocol by Iyer et al^47^. UMAP projections show **b**, protocol day; **c**, expression of PODXL / BNC or TCF21 / THY1 populations; **d**, PODXL or TCF21 positive lineage separation; and **e**, heart cell classification by the naïve Bayes classifier. **f** a time-course of hESC-EPI cell-type composition during differentiation. Box plots of mean gene expression of **g**, *in vivo* epicardial gene modules; and **h**, selected foetal specific genes in adult, foetal as well as hESC-EPI cells during differentiation.

We then calculated the expression of epicardial gene modules in the hESC-EPI cell and found that the expression of the regenerative foetal epicardial module C increased during differentiation until it was similar to *in vivo* foetal epicardium (Figure 5g). In contrast, the adult epicardial module H was absent in hESC-EPI cells (Figure 5g). We also noticed that the mesothelial gene module D was more highly expressed in the PODXL+ branch A while the expression of an EPDC gene module A appeared higher in the TCF21+ branch B. Encouragingly, we also observed a transiently high initial expression of the cell cycle associated gene module I (Figure 5g). Finally, we found that epicardial-specific but selectively-foetal genes were expressed in hESC-EPI cells increasing throughout differentiation including SLIT3, CXCL14, NRP2, SFRP2, and TFPI2, as well as epicardial-specific collagens, TNNT1, and RAMP1 found in module C (Figure 5h). This suggests a conserved role for these genes in epicardial function across both *in vivo* and *in vitro* environments and highlights *in vivo* pathways that hESC-EPI cells may re-activate when administered in hESC-CM grafts.

### A regenerative network waits for active epicardium

Lastly, we expanded the therapeutic context for reactivating adult epicardium or for administering hESC-EPI cells by creating a broader network of potential paracrine targets. Using StringDB, which includes semi-curated protein-protein interactions, we focused on connecting the epicardial secretomes with the membranomes of adult cardiomyocytes, endocardium, or endothelium (Figure 6). Contrasting slightly with the CellPhoneDB analysis, we found that the majority of edges connected foetal epicardium to endothelial cluster 1 (small vessel endothelial cells (Supplementary figure 3)). Among these interactions we identified SLIT3 in both *in vivo* and hESC-EPI interacting with ROBO4 on small vessel endothelial cells. We also find a large cluster centred around the previously discussed TGFB3 incorporating the highly precise marker of foetal epicardium BMP3. This cluster also contains collagens (COL11A1 and COL9A3) shared in both and *in vivo* and hESC-EPI. Endothelial and endocardial targets of this cluster were TGFBR2 and ENG (endoglin), a glycoprotein with high-affinity binding for TGFB3. Endothelial clusters also connected to epicardial NRP2, SEMA3B and SEMA3C through the membrane targets KDR, FLT1 and PLXNA2 forming a trio of angiogenic effectors. Finally, the foetal WNT signature WNT2B, RSPO1, SFRP2 / 5, and TFPI2 clustered together and modestly suggested a connection with NOTCH signalling in endothelial 2 (larger vessels) through JAG1. These findings conclude the results and describe a ready-and-waiting network of angiogenic signalling in adult hearts directed only by foetal epicardium, thus revealing the potential of restoring foetal epicardial states for human heart regeneration.

**Figure 6.**
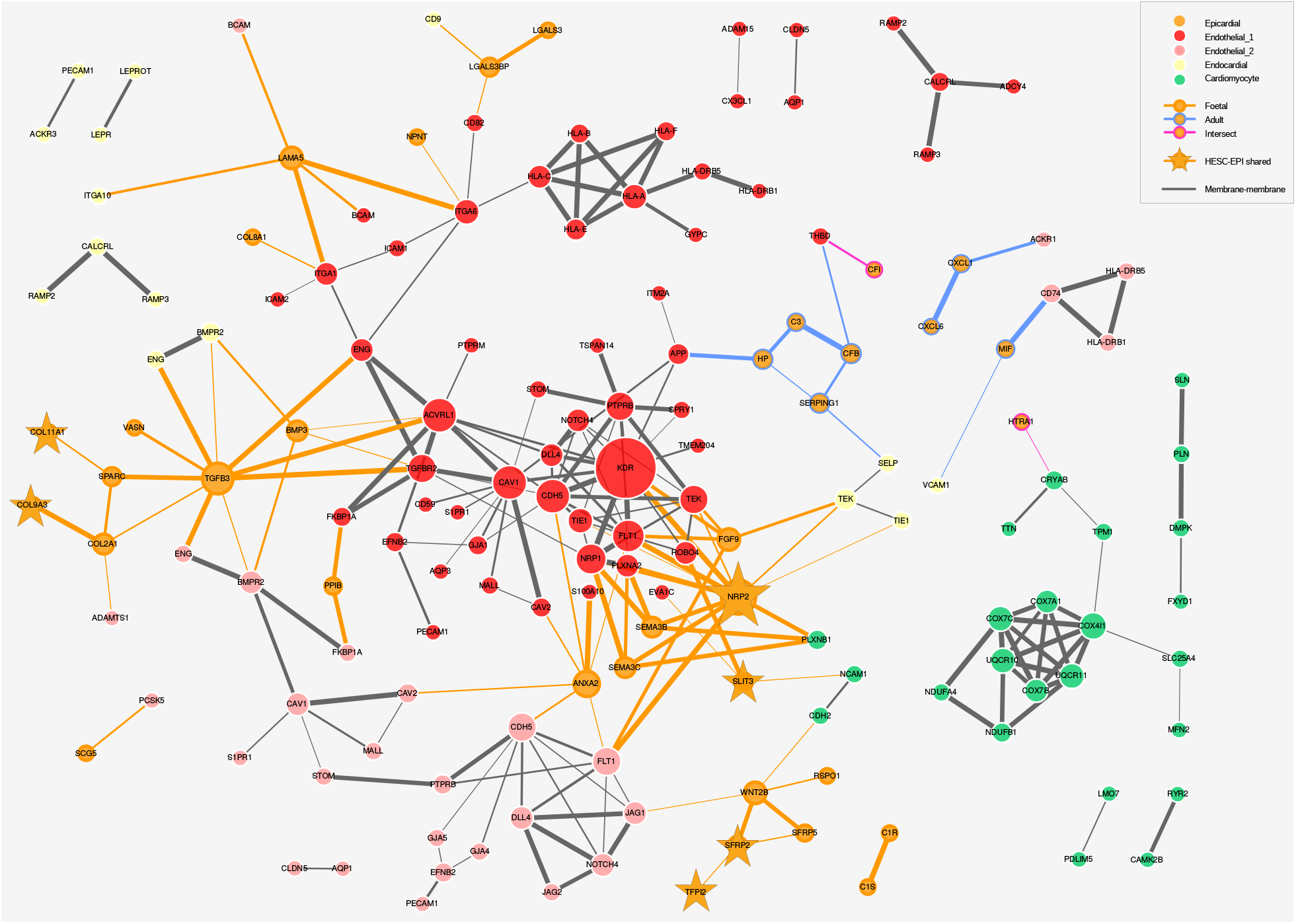
A broader network of potential interactions directed by secretions from both adult and foetal epicardium with fewer edges from adult nodes. Nodes were derived from differentially upregulated genes and filtered for secreted proteins in epicardium, or membrane-components in endothelial cells, endocardial cells or cardiomyocytes after differential expression analysis between cell clusters. Selected genes denoted by the star were also shared by hESC-EPI cells as cases of special interest. Grey edges illustrate potential membrane-component interactions and edge thickness denotes the overall score of the interaction according to stringDB while the size of each node is proportional to node degree.

## Discussion

Evidence suggests that epicardial activity is an important element of heart regeneration. On the one hand, active epicardium plays a substantial role in restoring cardiac function in adult zebrafish and newts as well as in developing mammalian systems. On the other hand, the epicardium is reportedly quiescent in adult mammalian and human hearts, which lack regenerative capabilities. However, despite the apparent importance of the epicardium, few studies have defined the age-associated changes in regenerative programs in human epicardial cells presenting an opportunity for finding novel therapeutic mechanisms in treating ischaemic injury. Addressing this unexplored space, we combined and compared foetal and adult hearts from humans at single-cell resolution for the first time and focused on epicardial cells within them. We revealed both compositional and molecular differences between the adult and foetal epicardium that, in part, may underpin lost regeneration seen in adult human hearts. We found that the adult epicardium may i) have a limited population of mesenchymal EPDCs, ii) have a lower volume of intercellular signalling, iii) lack specific regenerative and angiogenic gene programs found specifically in foetal epicardium and iv) be more primed for response to immune stimuli.

Our foetal-specific population of epicardial cells shared molecular characteristics of EPDCs such as those found in mouse and chick hearts^24,31^. However, in contrast with these studies, we found that our putative EPDC population was positive for the more mesothelial-associated LUM (lumican) but not expressing neonatal regeneration-associated proteoglycan AGRN (agrin). We may only have found these cells in foetal hearts as the adult epicardium may have a slower migratory response and may not infiltrate the myocardium^48^. Additionally, we made no effort to locate these cells experimentally and it remains unclear where they are situated within the myocardial tissue. That being said, our labelling of these cells as EPDCs was reasonable as their expression of epicardial and mesenchymal genes is indicative of EPDCs found within regenerating and developing myocardium, and our combination of clustering and classification strongly suggested an epicardial cell identity. To address the location of these cells and aid further research, we scored a library of epicardial markers in humans. This library was validated containing known epicardial genes and suggested that WT1 alone as used in some animals^27,49^ was not efficient in capturing all human epicardial cells. We suggest that researchers on human epicardium focus on the expression of precise epicardial markers shown here such as the ubiquitous UPK3B for all epicardial cells; SMPD3 and TNNT1 for foetal epicardial cells; and RBP4, HP, and PRR15L for adult epicardial cells. Additionally, the separation of mesothelial cells from putative mesenchymal cells may be achieved using SFRP5 in foetal states or ITLN1 and PRG4 across ages.

Our results agree with current understanding of adult epicardial quiescence. However, we demonstrated that this was directly related to absent epicardial-specific signals involved in angiogenesis, proliferation and survival. For human heart regeneration, one aim is to restore epicardial activity by reverting the adult epicardium to foetal states or by administering active epicardial cells generated from pluripotent stem cells^47,50^. Our study provides a road-map for this translational effort as for the first time we have revealed a transcriptome-wide division between adult and foetal epicardium which outlines the ingredients needed to bring foetal-like regenerative function back into adult epicardium. Firstly, the regenerative human epicardium may contain foetal angiogenic programs through NRP2, VEGFA, CXCL14 and SLIT3. These interactions have been confirmed in mice where epicardial SLIT2-mediated co-localisation with ROBO4 expressing endothelial cells was essential for proper vascularisation^31^. ROBO4 is further implicated in angiogenesis in human tissue^51^ and SLIT / ROBO signalling may also involve CXCL12 / CXCR4 interactions as provided by the epicardium^52,53^. These epicardial interactions with adult endothelial cells may be key as new vessel growth is likely sourced from pre-existing endothelial cells^54^. We find interest in CXCL14 as a highly-specific early foetal marker identified in our study as an allosteric modulator of CXCL12 / CXCR4, which may offer associated with developmental and regenerative coronary vasculogenesis. Furthermore, as CXCL14 is not found in animal studies of the epicardium, it may also be a key difference between animal and human epicardial signalling. Secondly, we should also aim to reactivate paracrine WNT signalling via SFRPs 2 and 5, and WNT2B. Previous animal studies showed high SFRP2 expression during cardiogenesis and regeneration with anti-fibrotic properties, specifically in post-injury epicardium^55–58^. However, SFRP5 has not been seen in animal studies and may be more relevant in humans. Studies found that SFRP5 was inversely proportional to cardiovascular disease risk factors, positively correlated with faster recovery after MI, and seen to protect against reperfusion injury^59–61^. Although the local targets of these WNT proteins may unknown, evidence suggests that WNT2B may increase proliferation in cardiomyocytes and fibroblasts^62,63^ and that restoring WNT signalling may key to adult heart regeneration. Other vital epicardial ingredients to include involve extracellular matrix remodelling, proliferation, and survival of myocardial tissue driven by TGFB3, BMP3, RSPO1^64^ and a variety of epicardial specific collagens such as COL11A1^65^. Lastly, we demonstrated that hESC-EPI cells contain many of these ingredients and have proven effectiveness in animal model grafts^22^ giving confidence in this recipe, and that bringing foetal programs back into adult epicardium is a viable strategy for adult human heart regeneration.

A surprising finding in our study was an enrichment of immune response genes in the adult epicardium. Experimental evidence in mice suggests that a rapid transient immune response increases regeneration^15^. While not disagreeing with this, our study may have revealed a novel shift in epicardial states and a persistent elevated responsiveness that may be intrinsic to normal human ageing. Additionally, we also found that immune cells were the most similar cell type between adult and foetal hearts implying an age-associated shift in epicardial sensitivity rather than a change in immune cell stimulation. This predicted change in epicardial immune response warrants further investigation as it might preclude heart regeneration in humans over time scales not usually covered in animal studies.

It is important to note that our study did not capture adult hearts from a diseased population but instead focussed on the healthy state. Therefore we could not compare the active foetal epicardium to injury-reactivated epicardium in post-MI adult samples. We consider that these foetal programs could also become expressed in the injury-reactivated adult epicardium. For example SFRP2 and SLIT3 are expressed in adult mice after injury^58^. However, one experiment in neonatal mice revealed an increase in RSPO1 in the regenerative P1 but not in non-regenerative P7 hearts after MI^16^. This independently validates our use of healthy-state adults as a model of non-regenerative epicardium in humans as RSPO1 was also decreased in our adult epicardium. An additional caveat of our study was that we compared epicardial cells most likely sourced from ventricular and apical regions in the foetus and atrial regions in the adult. This may have influenced the study as the composition of epicardium may be region dependent^1^. On a final note, it is incorrect to assume that the entire regenerative capacity of the heart rests upon the active epicardium; other cells also play a major role in regeneration. Addressing this comprehensively is beyond the scope of this study. However, our integrated dataset may also be used for future tissue-targeted and organ-wide studies on the age-associated changes in the human heart.

The next step for clinical translation is to disentangle the gene networks that regulate adult and foetal epicardial states. In doing so we might identify the molecular switches required to revert the adult epicardium back into a foetal state and restore these key pathways. Finally, because we have detailed both active and inactive states of the human epicardium, benchmarked cross-species epicardial markers in humans, and shown that stem cell-derived epicardium contains regenerative epicardial programs, this study serves as a valuable road map towards reactivating the adult epicardium and promoting heart regeneration in adult humans.

## Methods

### Collection of foetal cells for sequencing

Human foetuses were obtained following elective termination of pregnancy with full consent and stored overnight in Hibernate-A Medium (Gibco) at 4° C. The next day, the apex of the heart was dissected, and dissociated^66^. Briefly, tissue was dissociated using 6.6mg/ml Bacillus Licheniformis protease, 5 mM CaCl2 and 20U/ml DNase I, where the mixture was triturated on ice for 20 s every 5 minutes until clumps of tissue were no longer visible. The digestion was stopped with ice cold 10 % FBS in PBS. Cells were then washed with 10 % FBS, resuspended in 1 ml PBS and viability assessed using Trypan blue. Cells were submitted for 10x library preparation for 3’ single cell sequencing on a NovaSeq 6000 (Illumina) at the Cancer Research UK Cambridge Institute. Sample F5 was enriched for epicardium by carefully peeling the outermost cells from the myocardium and was sequenced separately at the Sanger Institute.

### hESC-EPI cell differentiation for the scRNAseq time course

Differentiation of epicardium was carried out according to our previously published protocols^47^. HESCs were initially differentiated into lateral plate mesoderm (LM) in the presence of FGF2 and BMP4. The LM is then exposed to WNT3a, BMP4 and retinoic acid resulting in hESC-EPI after 8 to 9 days. For this scRNAseq experiment we sampled days 1, 2, 3, 4, 8, and 9 of differentiation after the LM stage.

### RNA sequencing preprocessing

Cellranger version 3.02 with 10x v3 chemistry was used to demultiplex and generate matrices for downstream analyses of foetal samples F1, F2, F3, F4, F6 and F7 as well as the hESC-EPI samples. Default parameters were used and reads were aligned to the human reference genome hg38. The read count matrix for sample F5 was obtained using a previous version of Cellranger. Cleaning of the read count matrices was carried out in R removing all cells with a low and high number of UMIs (1,000 < # UMIs < 15,000), low number of genes expressed (# genes < 300), and a high fraction of UMIs found in mitochondrial genes (% MT > 15). These thresholds were chosen following the boundaries of the adult dataset and applied consistently across to reduce between donor effects. Doublets in foetal datasets were called using the software solo^67^ and were removed from the dataset after initial matrix pre-processing. Genes not expressed in any cell were removed from further analysis and the resulting matrices. Erythrocyte contamination in foetal samples was identified and removed using a single-variable two compartment gaussian mixture model (GMM) on the mean expression of haemoglobin genes. For this study, the hESC-EPI dataset was sampled taking only 300 cells from each time point. Cleaned UMI matrices were then compiled into Seurat objects for the majority of downstream analyses.

### Classification model

The heart cell classifier was constructed on the first collected and pilot sample F5 only. After quality control and matrix processing, the UMI counts of cells were transformed into feature frequency matrices by dividing counts by the total number of counts in each cell. We then generated a 50 dimensional uniform manifold approximation and projection (UMAP) and performed hierarchical density-based clustering using the algorithm HDBSCAN^68^ (Supplementary Figure 2a). A total of 12 different clusters were found and manually annotated, including a noise cluster of unknown cell type and a red blood cell cluster which were both associated more with lower quality cells (Supplementary Figure 2b). Cluster-based differential expression analysis using Wilcoxon rank-sum tests followed by a random forest algorithm was then used to rank gene features by their importance. A total of 647 genes were used to training the naïve Bayes classifier (NBC) (Supplementary Figure 2c). For the NBC, likelihood distributions for each gene were created using fixed-bandwidth kernel density estimates (KDEs) over the gene-frequency distributions within each cell type and the prior categorical distribution governing the probability of each cell type was determined by the fraction of cells belonging to each of the 12 clusters. Predictions on novel data were carried out by first transforming each sample’s gene counts matrices into gene frequencies. Then, posterior likelihood estimates of each gene-frequency value were calculated for each class using linear interpolation of adjacent points of KDE distributions described by the trained model. Finally, cells were assigned to cell type classes based on the maximum likelihood calculated by the product sum of posterior likelihoods and prior probabilities.

### Integration of adult and foetal data with stratified sampling

We used a stratified sampling strategy tailored to remove donor and cell-type composition biases within and between the samples and stages. Firstly, datasets were integrated within each stage separately and Louvain clustering was performed (Figure 1e & f). Epicardial clusters were identified by further subclustering an aggregate of epicardial, fibroblast, smooth muscle cell, and cycling cell clusters which were then collapsed together when the composition of HCC predictions within clusters were highly correlated (Supplementary Figure 3). Updating the initial clustering on foetal data resulted in 21 cell-type stratifications of foetal cell types (Supplementary Figure 3b & d). For adult data, we used the 47 cell type stratifications from the “cell_states” annotation previously assigned in the initial study^28^ (Supplementary Figure 3). The adult annotation set was deliberately chosen to include as many sub-clusterings of cell types as possible. Using these stratifications we further stratified the data by donor on each dataset. To do this we built an algorithm using a single parameter *x* to control the sampling rate across *K* clusters of size *N*_*K*_ where *x* is the approximate sample size to take from each cluster *K*. Iteratively, our algorithm randomly sampled cells from each donor in each cluster *K* without replacement until the number of cells in the newly sampled cluster, *k*, exceeded *x* (*n*_*k*_ > *x*).

For cases where *N_K_ < x*, the number of sampled cells is equal to the cluster size (*n*_*k*_ = *N*_*k*_) effectively sampling all available cells and donors. The effect of this was to spread the heart donors evenly within each cell type cluster with donor-bias, and reduce the cluster size in cell types where there was an abundance of cells (Supplementary Figure 3b & c). Being both the cell type of interest and one of the smallest compartments of the heart, we fixed *x* _*f*_ *_oetal_* to capture all foetal epicardium. We then used the total number of foetal cells after sampling to calibrate *x*_*adult*_ manually to obtain a similar number of cells in both stages. The cells sampled from all datasets were then integrated using Seurat’s standard integration pipeline across the donor category^30^. Highly variable genes (HVGs) were calculated in each sample before integration and combined, taking the unique intersection of HVGs for integration features across the datasets and for use in downstream differential expression analyses. Clustering across the integrated dataset was performed using the Louvain algorithm in Seurat and divided hierarchically at different resolutions offering different granularities for analysis (Supplementary Figure 3g-j).

### Parallel marker analysis and CellPhoneDB

In a separate workflow prior to stage integration, the within-stage integrated datasets were filtered to retain only cells kept in the stratified sampling stage and labelled by the 23 clusters found after stage integration. Differential expression analysis was then carried out using a Wilcoxon rank-sum test in a one cluster vs all approach. The precision (the fraction of cells expressing the gene that are epicardium), recall (the fraction of all epicardial cells expressing the gene) and F scores for the identified markers were calculated using the fraction of gene-positive cells that were either epicardial or non-epicardial within adult hearts for the case of adult markers, foetal hearts for the case of foetal markers, and hearts of both stages for the case of the intersection. The F score was calculated as the harmonic mean between recall and precision 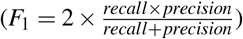. Log2 transformed counts acquired from the integrated matrix after stratified sampling were used as input into the statistical analysis pipeline from CellPhoneDB^69^.

### Binary data and gene-gene co-occurrence clustering for epicardial cells

An epicardial binary expression matrix was generated using a selected subset of genes defined by combining the top 1000 HVGs from the epicardial-only subset of cells and the epicardial-specific marker genes identified from the within-stage marker analysis. Gene count matrices were binarised by calling all counts above 0 as 1. We implemented the approach by Qiu, P^70^ in R to cluster gene-dropout patterns. Briefly, we calculated the co-occurrence of each gene with every other gene using chi-square statistics. Then, for each gene-pair, all chi-square statistics below a given threshold were discarded to retain only the high-scoring gene-gene pairs. The threshold to remove such low scoring gene-gene pairs was calculated using random permutations of the data. Then, gene pair chi-square statistics describing the gene dropout patterns were clustered using Louvain modularity clustering. In contrast with Qiu, P^70^, we performed clustering using the Rphenograph implementation of the Louvain clustering and chose to remove clusters with fewer than 20 genes. We consider this a hybrid approach because instead of using this method to broadly filter genes iteratively, we pre-filtered our gene set prior to co-occurrence clustering by the HVGs and epicardial-specific markers from the parallel marker analysis and viewed our gene set as either variable or important for epicardial cell transcriptional states. Cell clustering using this approach was then carried out using the mean values for each gene-module. This is considered equal to the fraction of genes expressed in each gene-module and used as a new matrix for Louvain clustering of cells. Gene sets from distinct epicardial gene expression modules were submitted to GprofileR using the R package gprofiler2^71^ using all filtered genes within our heart dataset as background.

### hESC-EPI differentiation time course and analysis

A sample of 300 cells was taken *in silico* from each time point and the cells were transformed into gene frequency and classified using the HCC. In parallel, we normalised, scaled and transformed the raw counts using Seurat and clustered the cells using Seurat’s implementation of Louvain clustering. The clusters were then labelled into lineages by their expression of PODXL or TCF21, after which the mean expression of the epicardial gene modules was calculated.

### Network Construction

Epicardial marker genes were filtered to retain only secreted protein gene-products while cardiomyocyte, endothelial 1 / 2, and endocardial marker genes were filtered to retain only membrane-bound gene products. Secreted or membrane-bound genes were listed on the human protein atlas (HPA). Gene nodes were then connected with edges described by their interactions in StringDB using an overall score threshold of above 400 and an experimental score of above 0 with the aim of capturing only experimentally authentic interactions. The network was visualised and analysed in Cytoscape^72^.

## Author contributions statement

V.K.S. conceived, designed and performed the analysis, and wrote the paper. H.D. coordinated and prepared foetal samples for sequencing, A.R. and L.V. contributed data sample F5, X.H. coordinated and collected foetal tissue samples, L.G. and S.S. conceived and designed the analysis, and wrote the paper. All authors reviewed and copy edited the manuscript.

## Acknowledgements

The authors would like to thank Roger Barker at the University of Cambridge for their assistance in obtaining the foetal tissue samples. This research was funded by the British Heart Foundation (BHF) Senior Fellowship [FS/18/46/33663] (SS, LG), Oxbridge BHF Centre for Regenerative Medicine [RM/17/2/33380] (VKS), and BHF grants [PG/17/24/32886] (LG) and [RG/17/5/32936] (HD). We also acknowledge core support from the Wellcome Trust and MRC to the Wellcome Trust – Medical Research Council Cambridge Stem Cell Institute. This research was funded in whole, or in part, by the Wellcome Trust [Grant Number: 203151/Z/16/Z]

## Additional information

### Data availability

The RNA sequencing data generated in this study will be submitted to the Gene Expression Omnibus (GEO) database or suitable repository and accession codes placed here.

