## Supplemental Figures for "Regenerative and non-regenerative transcriptional states of the human epicardium: from foetus to adult and back again"

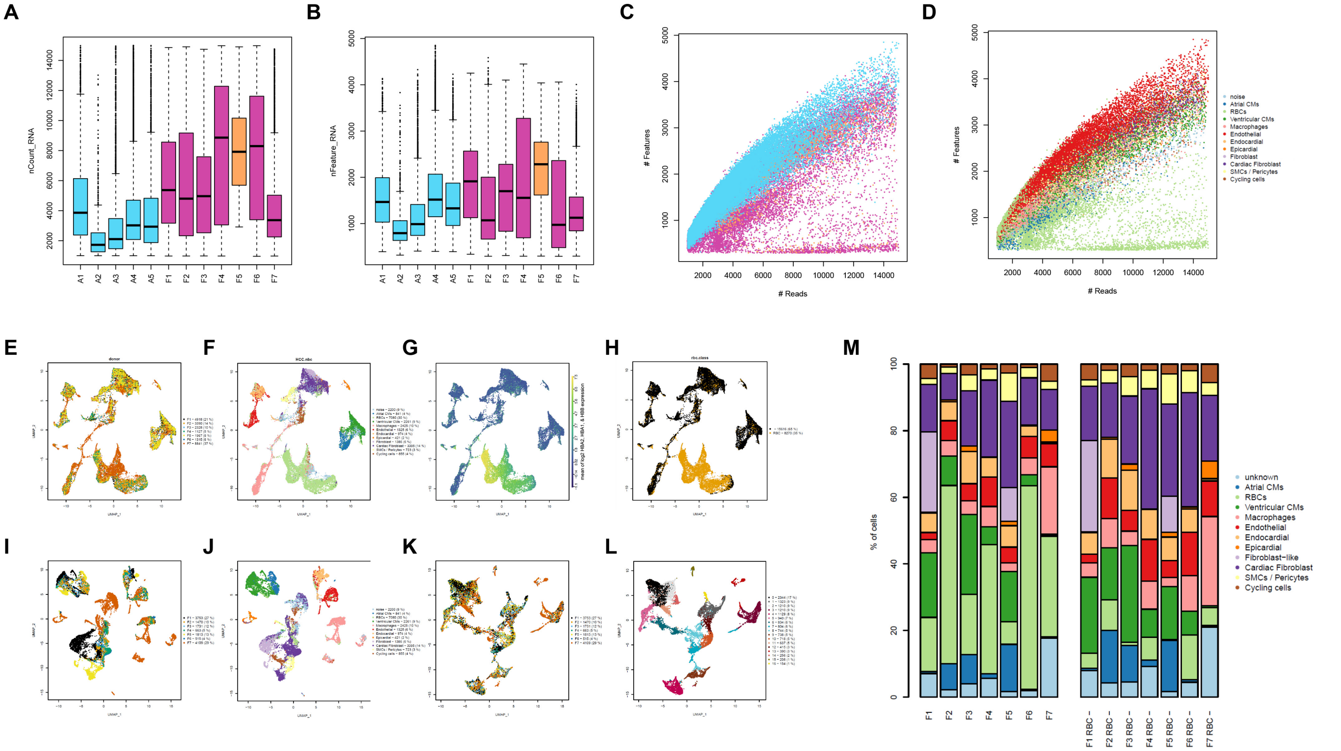


**Figure 1.** Quality control for all foetal samples after alignment, cleaning of erythrocyte contamination and identification of foetal clusters. **A**. UMI count distributions; **B**, feature count distributions; **C**, UMIs against features showing sample and; **D**, heart cell classification results; E-H, uniform manifold approximation and projections (UMAPs) of foetal samples before the removal of red blood cells (RBCs); **I-J**, UMAPs of foetal samples merged without batch correction and after integration (**K-L**); **M** heart cell classifier cell type distributions before and after RBC removal.

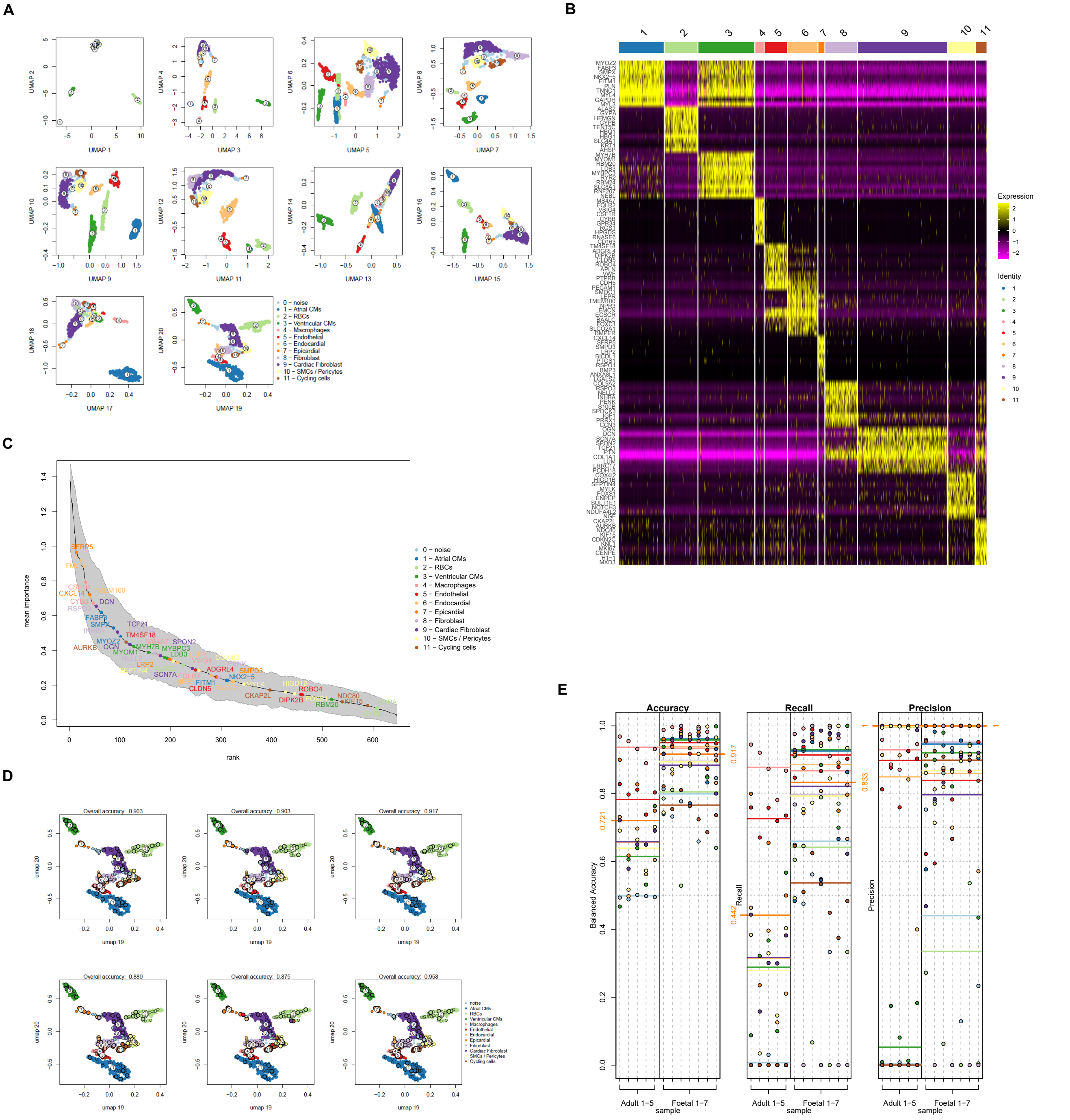


**Figure 2.** **A**. The first 20-components of a 50-dimensional uniform manifold approximation and projection (UMAP) of the gene-frequency matrix from sample F5 labelled by the 12 HDBSCAN clusters; **B**, top 10 gene markers for each HDBSCAN cluster identified using Wilcoxon ranked sum tests, of which the top 5 are located in; **C**, the ranking of gene features by the importance parameter from a random-forest feature optimization approach; **D**, UMAP components 19 and 20 with showing sampled cells belonging to each fold of cross-validation (small circle) as well as predictions (larger circle); and **E**, the accuracy, recall and precision of the classifier in all datasets determined independently by assessing the classifier's ability to capture cells belonging to putative clusters after within-stage integration of samples.

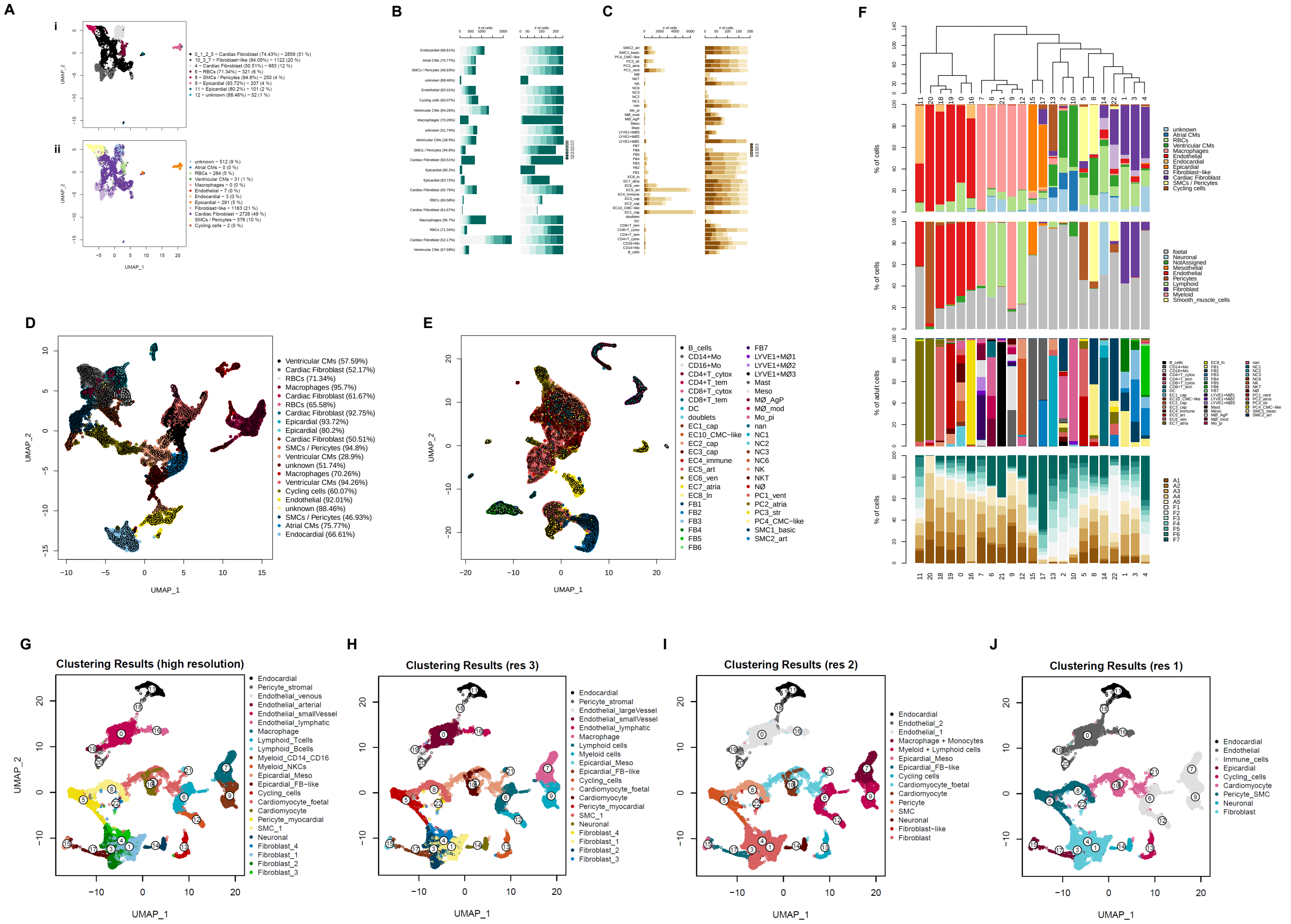


**Figure 3.** Stratified subsampling of datasets carried out by **A**, sub-clustering of epicardial, fibroblast and smooth muscle cell clusters identified in the foetal-integrated dataset; **B**, equally sampling of foetal clusters across donors and clusters to generate new equally-distributed foetal samples; **C**, equally sampling of adult "cell.states" across donors to generate new equally-distributed adult samples; **D**, umap of integrated foetal samples showing the locations of sub-sampled cells; **E**, umap of integrated adult samples showing locations of sub-sampled cells, **F**, highest resolution clusters for the integrated dataset after stratified sub-sampling ordered by hierarchical clustering of HCC-class compositions. Bar charts show a number of annotations types including HCC-class distribution, HCA-derived labels for adult cells (*cell.types* and *cell.states*) as a fraction of adult cells, and the distribution of the different samples among the clusters; **G-J**, decreasing resolutions of clustering in the integrated dataset after stratified sub-sampling.

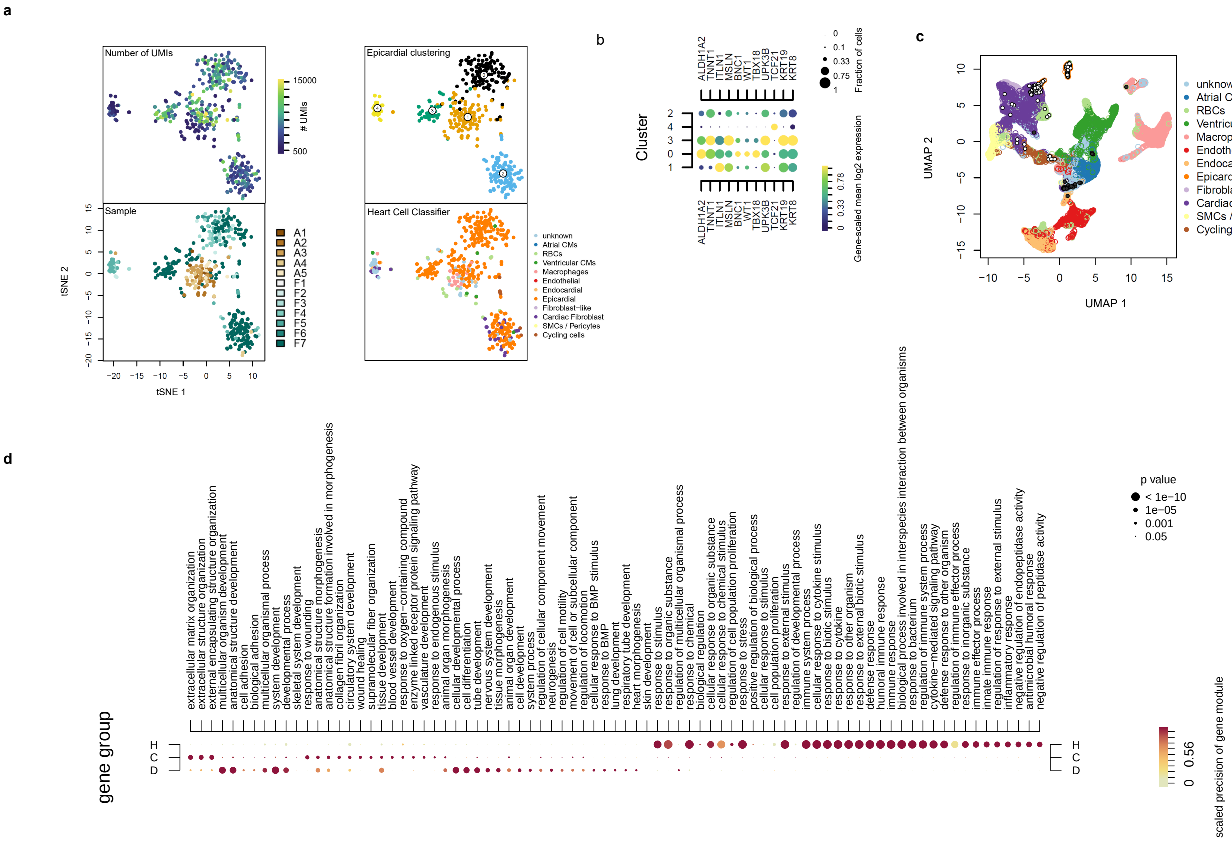


**Figure 4.** **a**, T-distributed stochastic neighbour embedded (tSNE) data of epicardial cells subset from the integrated dataset annotated with the library depth, heart cell classification, sample code, and sub-clustering; **b**, expression of epicardial genes across epicardial clusters; **c**, origin of both noisy (black) and non-noisy (white) epicardial cells on the dataset of all foetal samples; **d**, extended dot-plot showing additional significant terms within foetal specific gene modules C and D as well as adult cluster H.

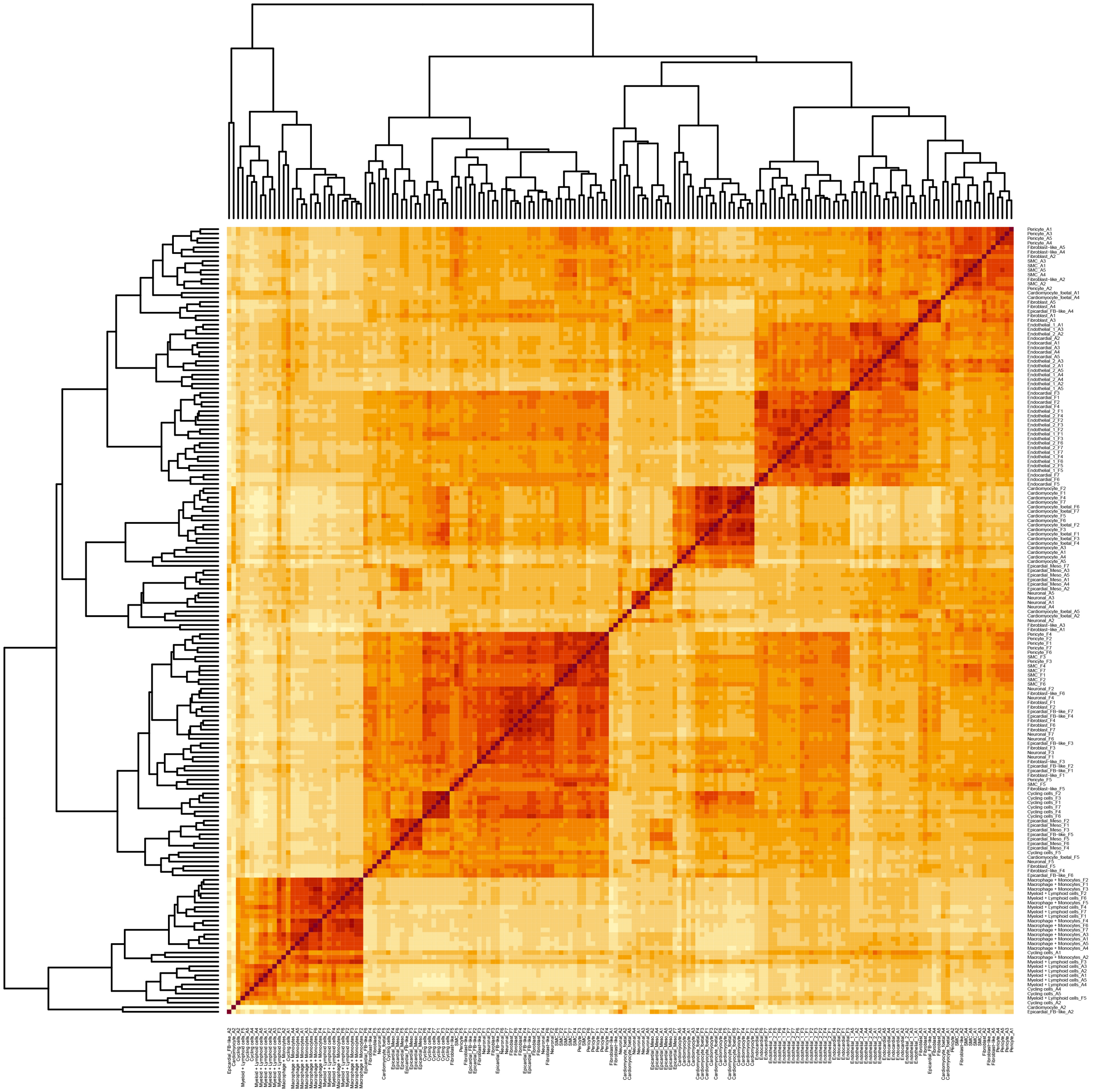

**Figure 5.** Pseudobulk downsampling of all cell types across adult and foetal donors and the results of hierarchical clustering.

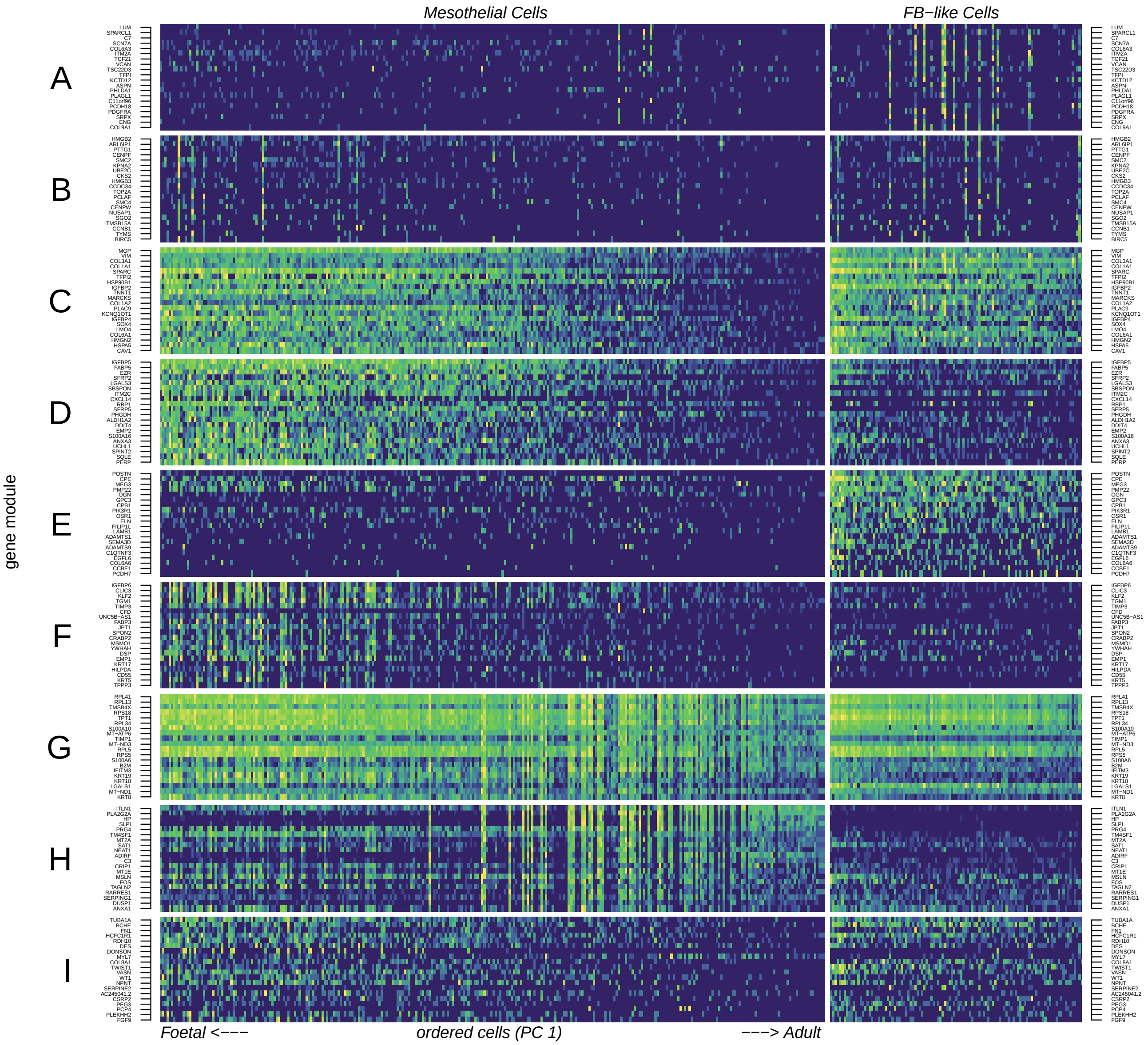


**Figure 6.** Gene module principal component analysis shows an age-associated switch in gene module commitment seen with ordering all epicardial cells by principal component 1 coordinates. The top 20 genes by expression were sampled from each module and shown here.
