## Supplemental Tables for "Regenerative and non-regenerative transcriptional states of the human epicardium: from foetus to adult and back again"

**Supplementary Table 1**. Sample information including sample labels, age, number of cells called by cellranger, and number of cells retained after quality control (QC) steps. Ages of foetal samples are approximate.

| Sample | Alt name | Age | # cells raw | # cells after QC |
|---|---|---|---|---|
| F1 | BRC2252 | 58 days | 7038 | 3753 |
| F2 | BRC2262 | 61 days | 5487 | 1470 |
| F3 | BRC2251 | 64 days | 3540 | 1731 |
| F4 | BRC2263 | 66 days | 2729 | 663 |
| F5 | alexsc | 67 days | 2486 | 1813 |
| F6 | BRC2256 | 69 days | 3265 | 515 |
| F7 | BRC2260 | 80 days | 10787 | 4109 |
| A1 | D3 | 55-60 years | 3856 | 3743 |
| A2 | D4 | 70-75 years | 1523 | 881 |
| A3 | D5 | 65-70 years | 6105 | 4721 |
| A4 | D6 | 65-70 years | 20889 | 19439 |
| A5 | D7 | 60-65 years | 5089 | 4318 |

**Supplementary Table 2.** Typical mean results from one 6-fold cross-validation run of the foetal sample F5. A sample of 36 cells was taken from each cluster of the total dataset in this run of 6-fold cross validation. Data points can be observed in the UMAP in Supplementary figure 2.

| Cell type | Recall | Precision | Balanced Accuracy |
|---|---|---|---|
| Unknown / Noise | 0.306 | NA | 0.645 |
| Atrial CMs | 1.000 | 1.000 | 1.000 |
| RBCs | 0.944 | 1.000 | 0.972 |
| Ventricular CMs | 1.000 | 1.000 | 1.000 |
| Immune cells | 0.972 | 0.976 | 0.985 |
| Endothelial | 0.944 | 0.935 | 0.968 |
| Endocardial | 0.944 | 0.925 | 0.968 |
| Epicardial | 1.000 | 0.976 | 0.999 |
| Fibroblast-like | 0.944 | 0.877 | 0.966 |
| Cardiac Fibroblast | 0.944 | 0.866 | 0.965 |
| SMCs / Pericytes | 0.917 | 0.858 | 0.951 |
| Cycling cells | 0.972 | 0.817 | 0.975 |

**Supplementary Table 3**. The top 100 differentially upregulated genes of epicardial cluster 15 compared with all other clusters identified using Wilcoxon rank-sum tests on the adult and foetal integrated dataset. Genes are ranked according to their log 2 fold change against the overall dataset excluding those with log2 fold-change lower than 0.5 or a p value greater than $1x10^{-10}$. Epicardial genes in only cluster 15 are shown in blue.

| Gene | Log2 FC | Adj pvalue | cluster % | dataset % | Epi_cluster |
|---|---|---|---|---|---|
| ITLN1 | 5.12 | 1.26E-158 | 100 | 50.4 | Both |
| HP | 5.11 | 5.06E-160 | 100 | 56.6 | Mesothelial |
| PRG4 | 3.78 | 1.90E-156 | 99.6 | 61.6 | Both |
| SLPI | 3.76 | 1.23E-158 | 100 | 69.3 | Both |
| PLA2G2A | 3.12 | 1.51E-150 | 100 | 70.4 | Both |
| KRT18 | 2.88 | 1.15E-154 | 99.6 | 50 | Both |
| KRT19 | 2.75 | 1.37E-159 | 99.2 | 41 | Both |
| KRT8 | 2.42 | 6.33E-153 | 99.2 | 47.4 | Both |
| UPK3B | 2.38 | 1.14E-165 | 99.2 | 28.8 | Both |
| TIMP1 | 2.15 | 1.58E-143 | 100 | 86.9 | Both |
| C3 | 2.14 | 2.15E-143 | 100 | 70.4 | Both |
| CCDC80 | 2.05 | 3.43E-137 | 99.6 | 66.5 | Both |
| SAT1 | 2.02 | 2.82E-133 | 100 | 90.7 | Mesothelial |
| SGK1 | 1.80 | 3.77E-148 | 100 | 75 | Both |
| CYSTM1 | 1.72 | 3.30E-139 | 100 | 85.2 | Both |
| HAS1 | 1.54 | 1.02E-158 | 99.6 | 54.5 | Both |
| SYT4 | 1.52 | 1.55E-176 | 99.2 | 26.6 | Both |
| RARRES2 | 1.46 | 5.36E-126 | 99.2 | 68.6 | Both |
| INMT | 1.45 | 1.76E-129 | 99.6 | 77.3 | Both |
| WFDC2 | 1.45 | 4.56E-156 | 99.6 | 55.5 | Mesothelial |
| SLC39A8 | 1.45 | 1.00E-149 | 99.6 | 66.3 | Both |
| FOS | 1.43 | 2.21E-109 | 100 | 86 | Mesothelial |
| TM4SF1 | 1.41 | 8.44E-107 | 100 | 80.8 | Mesothelial |
| CALB2 | 1.39 | 8.55E-144 | 96 | 30.4 | Both |
| EFEMP1 | 1.38 | 1.35E-118 | 99.6 | 71.5 | Both |
| MSLN | 1.37 | 2.53E-138 | 95.2 | 35.6 | Both |
| CFI | 1.36 | 1.30E-130 | 98.4 | 78.1 | Both |
| NNMT | 1.35 | 1.76E-114 | 99.6 | 78.5 | Both |
| UPK1B | 1.34 | 4.18E-150 | 98 | 37.9 | Both |
| AQP1 | 1.32 | 3.66E-104 | 100 | 84.7 | Mesothelial |
| CEBPB | 1.32 | 1.25E-116 | 100 | 87.2 | Mesothelial |
| PROCR | 1.30 | 8.97E-114 | 96.8 | 73.4 | Both |
| CXADR | 1.29 | 1.32E-149 | 98.4 | 46.6 | Both |
| CLDN15 | 1.27 | 7.75E-142 | 98.4 | 59.1 | Both |
| CHI3L1 | 1.24 | 4.12E-142 | 96.8 | 56.6 | Both |
| LINC01133 | 1.22 | 1.26E-140 | 97.6 | 46.8 | Both |
| MGST1 | 1.21 | 2.23E-98 | 100 | 76.8 | Both |
| S100A10 | 1.16 | 1.11E-119 | 100 | 94.5 | Both |
| PTGIS | 1.14 | 1.31E-123 | 99.6 | 67.5 | Both |
| FLRT3 | 1.09 | 2.16E-155 | 99.2 | 54.2 | Both |
| TGM1 | 1.06 | 2.63E-143 | 96.4 | 38.9 | Both |
| SELENBP1 | 1.05 | 5.14E-117 | 99.6 | 67.3 | Mesothelial |
| RPL22L1 | 1.04 | 3.02E-101 | 99.2 | 76.6 | Mesothelial |
| ZFAS1 | 1.04 | 6.86E-99 | 100 | 86.3 | Mesothelial |
| PDPN | 1.03 | 1.64E-137 | 99.2 | 60.4 | Both |
| GCHFR | 1.03 | 3.34E-134 | 98.8 | 73.9 | Mesothelial |
| CD200 | 1.03 | 1.71E-141 | 98.8 | 62 | Mesothelial |
| RARRES1 | 1.02 | 6.03E-86 | 98.4 | 70.8 | Both |
| ID2 | 1.02 | 1.38E-90 | 99.2 | 86.4 | Mesothelial |
| CFH | 1.01 | 2.54E-94 | 100 | 73.7 | Both |

| | | | | | |
|---|---|---|---|---|---|
| KLF6 | 1.00 | 2.63E-100 | 100 | 93.1 | Mesothelial |
| HILPDA | 0.98 | 1.32E-126 | 97.2 | 65.3 | Both |
| ALOX15 | 0.98 | 1.01E-173 | 98.8 | 29 | Mesothelial |
| MT1E | 0.96 | 1.20E-92 | 98.8 | 82.5 | Both |
| TPT1 | 0.94 | 2.73E-127 | 100 | 99.6 | Mesothelial |
| PTPRF | 0.93 | 5.40E-118 | 93.6 | 51.7 | Mesothelial |
| DUSP1 | 0.93 | 6.11E-81 | 100 | 94.4 | Mesothelial |
| ERRFI1 | 0.89 | 2.07E-139 | 98.4 | 76.5 | Mesothelial |
| PHYHIP | 0.88 | 8.89E-140 | 98 | 65.1 | Both |
| LAMA4 | 0.87 | 7.58E-109 | 98.8 | 65.7 | Mesothelial |
| PRR15 | 0.86 | 6.99E-156 | 98.4 | 46 | Mesothelial |
| RPS8 | 0.85 | 6.11E-128 | 100 | 99.1 | Mesothelial |
| EPHB6 | 0.85 | 9.80E-104 | 92 | 66.8 | Both |
| SNCA | 0.83 | 4.52E-124 | 98 | 52.7 | Mesothelial |
| KRT7 | 0.83 | 2.25E-82 | 86.5 | 58.6 | Mesothelial |
| RBP4 | 0.82 | 1.77E-112 | 91.2 | 38.5 | Mesothelial |
| TMEM176A | 0.82 | 1.79E-78 | 93.2 | 74.9 | Both |
| LDHA | 0.82 | 1.37E-70 | 99.6 | 93 | Both |
| DMKN | 0.81 | 7.83E-88 | 89.2 | 53.2 | Mesothelial |
| RPS12 | 0.78 | 8.67E-113 | 100 | 99.2 | Mesothelial |
| JUN | 0.76 | 8.52E-52 | 98.8 | 88.9 | Mesothelial |
| RHOB | 0.74 | 6.86E-68 | 100 | 91.1 | Mesothelial |
| PLIN2 | 0.74 | 9.88E-101 | 98.8 | 77.2 | Both |
| KCNT2 | 0.74 | 2.68E-145 | 98.4 | 64.2 | Mesothelial |
| SLC16A1 | 0.73 | 1.11E-110 | 95.6 | 68.8 | Mesothelial |
| RPL12 | 0.72 | 2.97E-109 | 100 | 98.7 | Mesothelial |
| MEDAG | 0.71 | 2.82E-85 | 91.2 | 69.5 | Both |
| PDLIM4 | 0.71 | 1.25E-116 | 98 | 53.5 | Mesothelial |
| MT2A | 0.71 | 1.03E-61 | 99.2 | 92.5 | Mesothelial |
| GFPT2 | 0.71 | 5.52E-111 | 97.2 | 61.6 | Both |
| SLC4A4 | 0.70 | 2.33E-139 | 98.4 | 55.6 | Mesothelial |
| LY6E | 0.68 | 3.92E-57 | 96.4 | 88.6 | Mesothelial |
| PIM1 | 0.67 | 7.41E-125 | 98.8 | 69.2 | Mesothelial |
| NDN | 0.67 | 4.78E-72 | 96 | 62.6 | Mesothelial |
| MYRF | 0.67 | 5.80E-70 | 82.9 | 39.1 | Mesothelial |
| RPS4X | 0.67 | 4.22E-103 | 100 | 98.1 | Mesothelial |
| VSIG2 | 0.67 | 4.32E-153 | 98.8 | 56.5 | Mesothelial |
| COBLL1 | 0.67 | 3.57E-88 | 98.4 | 80.3 | Mesothelial |
| CTNNB1 | 0.66 | 9.37E-47 | 98.4 | 86.7 | Mesothelial |
| RPL7A | 0.66 | 2.30E-84 | 100 | 98.8 | Mesothelial |
| PRDM6 | 0.65 | 4.86E-153 | 98.4 | 45 | Mesothelial |
| CDH3 | 0.65 | 6.75E-131 | 94 | 37.5 | Mesothelial |
| PKHD1L1 | 0.65 | 7.75E-121 | 97.6 | 64.8 | Mesothelial |
| RPL3 | 0.64 | 1.36E-88 | 100 | 98.9 | Mesothelial |
| MT-ND4 | 0.63 | 1.46E-92 | 100 | 100 | Mesothelial |
| ZNF593 | 0.63 | 1.45E-97 | 98.8 | 77.1 | Mesothelial |
| SERPING1 | 0.62 | 4.07E-54 | 99.2 | 86.6 | Both |
| NAMPT | 0.62 | 9.27E-104 | 100 | 88.2 | Mesothelial |
| RBP1 | 0.61 | 2.33E-48 | 92 | 69.3 | Both |
| ATP1B1 | 0.61 | 2.25E-69 | 96 | 72.4 | Mesothelial |

**Supplementary Table 4**. The top 88 differentially upregulated genes of epicardial cluster 17 compared with all other clusters identified using Wilcoxon rank sum tests on the adult and foetal integrated dataset. Genes are ranked according to their log 2 fold change against the overall dataset excluding those with log2 fold-change lower than 0.5 or a p value greater than $1x10^{-10}$. Epicardial genes in only cluster 17 are shown in red.

| Gene | Log2 FC | Adj pvalue | cluster % | dataset % | Epi_cluster |
|---|---|---|---|---|---|
| PLA2G2A | 0.85 | 1.06E-65 | 100 | 70.6 | Both |
| CCDC80 | 0.84 | 5.77E-60 | 97.9 | 66.8 | Both |
| TIMP1 | 0.83 | 6.52E-44 | 100 | 87 | Both |
| C3 | 0.73 | 2.02E-58 | 99.5 | 70.6 | Both |
| EFEMP1 | 0.65 | 1.48E-52 | 99.5 | 71.7 | Both |
| SLPI | 0.63 | 2.60E-98 | 98.9 | 69.5 | Both |
| NNMT | 0.60 | 1.23E-42 | 98.4 | 78.7 | Both |
| MGST1 | 0.59 | 5.26E-46 | 100 | 76.9 | Both |
| C1R | 0.56 | 2.23E-33 | 98.4 | 77 | FB-like |
| RARRES1 | 0.55 | 5.42E-49 | 98.9 | 70.9 | Both |
| KRT19 | 0.53 | 1.39E-70 | 89.9 | 41.5 | Both |
| CFH | 0.51 | 2.57E-44 | 100 | 73.8 | Both |
| PTGIS | 0.51 | 4.36E-50 | 98.4 | 67.7 | Both |
| RARRES2 | 0.48 | 4.25E-46 | 96.8 | 68.8 | Both |
| LDHA | 0.47 | 1.89E-21 | 100 | 93 | Both |
| KRT18 | 0.46 | 2.37E-35 | 84.1 | 50.5 | Both |
| MT1X | 0.45 | 1.77E-21 | 99.5 | 87.6 | FB-like |
| S100A6 | 0.44 | 3.88E-22 | 100 | 98.3 | Both |
| C1S | 0.44 | 3.50E-34 | 98.4 | 76.1 | FB-like |
| KRT8 | 0.43 | 2.12E-26 | 77.8 | 48.1 | Both |
| CYSTM1 | 0.42 | 2.86E-30 | 98.4 | 85.3 | Both |
| LGALS3BP | 0.42 | 9.21E-20 | 93.1 | 75.3 | Both |
| ITLN1 | 0.41 | 3.30E-84 | 95.8 | 50.7 | Both |
| MFAP4 | 0.41 | 2.92E-36 | 92.6 | 57.6 | FB-like |
| IGFBP6 | 0.41 | 2.88E-39 | 96.3 | 70.8 | FB-like |
| LINC01133 | 0.40 | 1.67E-37 | 81.5 | 47.4 | Both |
| CHI3L1 | 0.40 | 2.82E-93 | 97.4 | 56.8 | Both |
| COL6A2 | 0.40 | 7.74E-21 | 98.9 | 84.5 | FB-like |
| SERPING1 | 0.39 | 9.71E-22 | 98.9 | 86.7 | Both |
| FSTL1 | 0.39 | 1.10E-23 | 93.1 | 73.9 | FB-like |
| UPK3B | 0.39 | 1.70E-40 | 78.8 | 29.6 | Both |
| NUPR1 | 0.39 | 2.85E-21 | 97.9 | 81.8 | Both |
| CLDN15 | 0.38 | 1.04E-34 | 82 | 59.6 | Both |
| MEDAG | 0.38 | 2.46E-20 | 79.4 | 69.8 | Both |
| HTRA1 | 0.37 | 1.86E-21 | 93.1 | 77.7 | Both |
| ID4 | 0.37 | 3.54E-25 | 92.6 | 71.8 | Both |
| PRG4 | 0.37 | 1.25E-91 | 97.4 | 61.8 | Both |
| PLAC9 | 0.37 | 5.17E-17 | 98.4 | 82.3 | FB-like |
| INMT | 0.36 | 2.15E-51 | 97.4 | 77.4 | Both |
| S100A10 | 0.36 | 5.23E-17 | 99.5 | 94.6 | Both |
| MSLN | 0.35 | 9.16E-37 | 78.8 | 36.2 | Both |
| CXADR | 0.34 | 6.62E-25 | 74.6 | 47.3 | Both |
| CST3 | 0.34 | 5.89E-12 | 99.5 | 94.6 | FB-like |
| HILPDA | 0.33 | 4.50E-35 | 91.5 | 65.6 | Both |
| SGK1 | 0.33 | 5.50E-36 | 95.8 | 75.2 | Both |
| RPLP0 | 0.33 | 2.02E-21 | 99.5 | 96.7 | Both |
| ARL4D | 0.33 | 1.37E-26 | 88.4 | 69.3 | Both |
| FLRT3 | 0.33 | 1.88E-33 | 78.8 | 54.8 | Both |
| COL1A2 | 0.33 | 2.87E-20 | 94.2 | 66.1 | FB-like |

| | | | | | |
|---|---|---|---|---|---|
| FBLN2 | 0.33 | 1.81E-27 | 95.8 | 77.8 | FB-like |
| SYT4 | 0.33 | 8.79E-101 | 94.2 | 27.1 | Both |
| RBP1 | 0.30 | 3.59E-22 | 90.5 | 69.5 | Both |
| UAP1 | 0.30 | 3.45E-34 | 95.2 | 71.5 | FB-like |
| TMEM176A | 0.30 | 1.22E-27 | 93.1 | 75 | Both |
| TGM1 | 0.29 | 2.21E-27 | 74.6 | 39.6 | Both |
| PROCR | 0.29 | 2.76E-17 | 85.7 | 73.7 | Both |
| CYBRD1 | 0.29 | 4.95E-31 | 97.4 | 78.5 | Both |
| COL1A1 | 0.29 | 2.85E-20 | 85.7 | 53.6 | FB-like |
| PLIN2 | 0.29 | 9.68E-32 | 96.8 | 77.3 | Both |
| GFPT2 | 0.29 | 2.07E-57 | 97.4 | 61.9 | Both |
| COL6A1 | 0.29 | 8.09E-18 | 94.7 | 75.9 | FB-like |
| MT1E | 0.29 | 1.27E-20 | 97.9 | 82.6 | Both |
| FBLN1 | 0.29 | 1.01E-37 | 97.9 | 60.9 | FB-like |
| DPT | 0.29 | 5.15E-44 | 98.4 | 74.2 | FB-like |
| PKDCC | 0.28 | 7.23E-35 | 91.5 | 66.9 | Both |
| OGN | 0.28 | 1.89E-13 | 80.4 | 62.5 | FB-like |
| PHYHIP | 0.28 | 1.06E-62 | 91.5 | 65.4 | Both |
| CFI | 0.28 | 1.41E-34 | 90.5 | 78.4 | Both |
| HAS1 | 0.28 | 3.10E-32 | 79.4 | 55.1 | Both |
| UPK1B | 0.28 | 1.32E-82 | 92.6 | 38.3 | Both |
| CALB2 | 0.28 | 4.88E-17 | 69.3 | 31.2 | Both |
| AEBP1 | 0.27 | 5.27E-25 | 93.1 | 75.5 | FB-like |
| PCOLCE2 | 0.27 | 3.81E-46 | 97.9 | 68.5 | FB-like |
| SLC39A8 | 0.27 | 5.95E-41 | 87.8 | 66.7 | Both |
| GXYLT2 | 0.27 | 2.07E-42 | 88.9 | 55.3 | Both |
| PTN | 0.27 | 6.03E-16 | 85.2 | 65.4 | FB-like |
| EPHB6 | 0.27 | 1.03E-15 | 73.5 | 67.3 | Both |
| RERG | 0.27 | 1.56E-37 | 95.8 | 67.6 | Both |
| FBN1 | 0.26 | 1.49E-37 | 98.4 | 68.2 | FB-like |
| ADGRD1 | 0.26 | 1.27E-40 | 94.7 | 73.2 | Both |
| TNXB | 0.26 | 1.12E-28 | 97.9 | 81.7 | FB-like |
| FN1 | 0.26 | 2.10E-23 | 98.9 | 78.8 | FB-like |
| VASN | 0.26 | 3.16E-17 | 87.3 | 69 | FB-like |
| ACKR3 | 0.26 | 3.48E-22 | 96.8 | 79 | FB-like |
| GPC3 | 0.26 | 1.74E-15 | 75.7 | 49.5 | FB-like |
| GAS1 | 0.25 | 7.03E-14 | 83.1 | 73.7 | FB-like |
| TCF21 | 0.25 | 1.51E-30 | 90.5 | 60.2 | FB-like |
| PDPN | 0.25 | 3.72E-14 | 75.1 | 61 | Both |

**Additional table legends (actual tables are in located separate excel sheets for ease of sharing).**

**Supplementary Tables 5a to i.** Listed gene module genes and the fraction of cells in each of the seven transcriptional states expressing each gene.

**Supplementary Tables 6a to c**. Raw expressions of Foetal, Intersect and Adult markers respectively in either foetal or adult mesothelial epicardial cell clusters (15) after parallel differential expression analysis using Wilcoxon rank-sum tests between clusters. Foetal and Adult sub-tables include log fold and p value columns of these markers within Foetal and Adult epicardium respectively for reference.
